# Structural characterization and dynamics of AdhE ultrastructures from *Clostridium thermocellum:* A containment strategy for toxic intermediates

**DOI:** 10.1101/2024.02.16.580662

**Authors:** Samantha J. Ziegler, Brandon C. Knott, Josephine N. Gruber, Neal N. Hengge, Qi Xu, Daniel G. Olson, Eduardo E. Romero, Lydia M. Joubert, Yannick J. Bomble

## Abstract

*Clostridium thermocellum*, a cellulolytic thermophilic anaerobe, is considered by many to be a prime candidate for the realization of consolidated bioprocessing (CBP) and is known as an industry standard for biofuel production. *C. thermocellum* is among the best biomass degraders identified to date in nature and produces ethanol as one of its main products. Many studies have helped increase ethanol titers in this microbe, however ethanol production using *C. thermocellum* is still not economically viable. Therefore, a better understanding of its ethanol synthesis pathway is required. The main pathway for ethanol production in *C. thermocellum* involves the bifunctional aldehyde-alcohol dehydrogenase (AdhE). To better understand the function of the *C. thermocellum* AdhE, we used cryo-electron microscopy (cryo-EM) to obtain a 3.28 Å structure of the AdhE complex. This high-resolution structure, in combination with molecular dynamics simulations, provides insight into the substrate channeling of the toxic intermediate acetaldehyde, indicates the potential role of *C. thermocellum* AdhE to regulate activity and cofactor pools, and establishes a basis for future engineering studies. The containment strategy found in this enzyme offers a template that could be replicated in other systems where toxic intermediates need to be sequestered to increase the production of valuable biochemicals.

## Introduction

Lignocellulosic biomass is one of the most attractive substrates for sustainable production of second-generation biofuels and other bioproducts (*1-5*). Consolidated bioprocessing (CBP) is emerging as a promising process to reduce costs by combining biomass solubilization and fermentation in one step without added enzymes (*6-9*). Due to its ability to quickly solubilize and utilize cellulose, *Clostridium thermocellum* is an ideal candidate organism for CBP and has previously been engineered to produce ethanol at high yield (*10, 11*). However, ethanol production using *C. thermocellum* is not yet an economical process due to limited titers (*12-14*). The main pathway for ethanol production in *C. thermocellum* involves the bifunctional aldehyde-alcohol dehydrogenase (AdhE), a gene that is often modified in strains selected for alcohol tolerance and other selective pressures, indicating its crucial role in *C. thermocellum* metabolism (*14-17*). AdhE functions in the anaerobic fermentation pathway and contains two domains: an aldehyde dehydrogenase domain (ALDH) for the reduction of acetyl-CoA to acetaldehyde and an alcohol dehydrogenase domain (ADH) to reduce acetaldehyde to ethanol. Both reduction processes natively use NADH as a cofactor and are reversible in the presence of NAD^+^ as a cofactor. Because AdhE is part of the native ethanol production pathway in many anaerobic bacteria, it is a key target protein in many biofuels and biochemicals-related studies (*14, 15, 17-20*).

Structurally, AdhE is an intriguing protein as it forms large, filamentous helical ultrastructures termed spirosomes, that could be necessary for function (*21, 22*). In the case of *E. coli* AdhE, spirosome formation seems vital for the forward reaction of acetyl-CoA to ethanol, while the reverse reaction is not impacted by the disruption of the ultrastructures (*23-25*). Prior structural studies on the *E. coli* AdhE complex captured both the extended and compact forms of the spirosome, with some biochemical evidence that the extended form was required for the reduction of acetyl-CoA to ethanol (*23, 24, 26*). Because *C. thermocellum* AdhE and *E. coli* AdhE have 62% sequence identity, we first aimed to determine the similarity between the AdhE from *C. thermocellum* and *E. coli*. To further understand the AdhE from *C. thermocellum*, we used cryo-electron microscopy (cryo-EM) to capture the structure of the spirosome ultrastructure in both its extended and compact forms. We further analyzed the results of our 3.28 Å extended *C. thermocellum* AdhE in combination with channel prediction software and molecular dynamics simulations. We determined that the native state of the *C. thermocellum* spirosome is extended (different from that of other structurally characterized AdhEs) and contains an enclosed channel between the ALDH and ADH active sites. In modeling the compact spirosome, we found that this channel leaks acetaldehyde to the bulk phase, which indicates that the extended spirosome might be crucial for toxic aldehyde intermediate channeling.

## Results and Discussions

### Spirosome conformation is determined by intrinsic sequence and local environment, but not by expression host

A fundamental aspect of this work was determining if the local environment of AdhE affected the formation of the spirosome and its conformation (extended *vs*. compact). This was especially important given that our structural studies utilized *C. thermocellum* AdhE heterologously expressed and purified from *E. coli*. Based on negative stain TEM data shown in Figure 1A, we found that spirosomes maintained the same conformation between endogenous and exogenous expression – both primarily extended in the case of *C. thermocellum* which represents the majority *apo* conformation of the *C. thermocellum* spirosome. Intriguingly, the extended conformation of the *C. thermocellum* spirosome remained unchanged when purified from *E. coli*, even though the native *apo E. coli* AdhE is purified primarily in the compact form (Figure 1B). Since Kessler *et al*. found that the addition of both NAD^+^ and FeSO_4_ was necessary to transform *E. coli* spirosomes from their compact to their extended forms, we investigated the effects of different reactants that could contribute to the inverse process in the purified, extended *C. thermocellum* spirosome (*21*). While our results were not as extreme as those of Kessler *et al*., we found that the addition of the forward reactants (NADH, acetyl-CoA, FeSO_4_) led to the compaction of spirosomes. Conversely, the addition of the reverse reactants (NAD^+^, ethanol, CoA, FeSO_4_) to the compact *E. coli* spirosomes resulted in more extended forms. To isolate which substrate was causing this shift, we made negative stained grids of the *C. thermocellum* spirosomes in the presence of each reactant and found that NADH and NAD^+^ were the major determining factors in the conformation of the spirosome (Figure 1B, Supplemental Figure 1). We also found that the expression host does not impact conformation of the spirosomes but does significantly affect their length (Figure 1C). *C. thermocellum* AdhE expressed in *E. coli* formed spirosomes that were, on average, approximately 20 nanometers shorter than those found in a native *C. thermocellum* lysate. This equates to spirosomes that contain, on average, approximately five to ten fewer AdhE monomers in the extended spirosome. The shorter average length was due largely to the lack of spirosomes that were longer than 100 nanometers in the exogenous expression (Figure 1C). This phenomenon has been observed previously with the exogenous expression of *V. cholerae* spirosomes in *E. coli*, which also resulted in shorter spirosomes (*27*).

**Figure 1:**
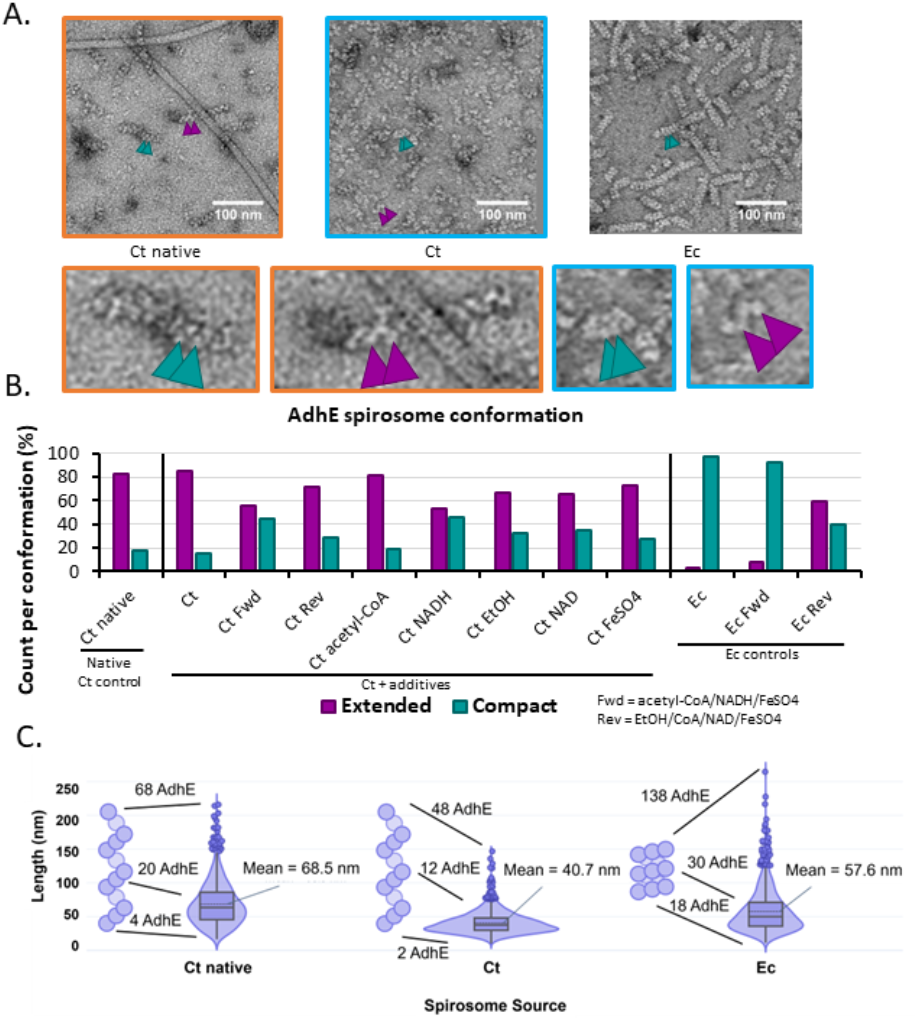
Negative-stain data of AdhE spirosomes indicate that conformation differs between bacteria. **A)** Sample negative stained images. Ct = *C. thermocellum*, Ec = *E. coli*. Compact spirosome indicated by teal arrows, extended by purple arrows. Zoom panels correlate by color. **B)** Measurement of spirosome conformation as determined in at least 500 instances. **C)** Violin plot of the lengths of spirosomes measured in 1250 instances – example spirosomes to the left of each group represents the majority conformation in the *apo* state. The difference in mean length was statistically significant for all three samples.

### High-resolution structure of the extended *C. thermocellum* AdhE spirosome

To understand why *C. thermocellum* AdhE is found in the extended spirosome form and *E. coli* AdhE in the compact form, we turned to cryo-EM to obtain a high-resolution structure of the *C. thermocellum* protein complex to analyze its interfaces as compared to previously published *E. coli* spirosome structures at the molecular level (*23, 24, 26*). Using standard cryo-EM techniques (see Materials and Methods), we were able to determine a 3.28 Å cryo-EM structure of the extended *C. thermocellum* AdhE spirosome (Figure 2). We compared multiple aspects of the protein complex to the previously published *E. coli* spirosome structures, including the interaction interfaces, predicted catalytic active sites, cofactor binding pockets, and potential channels that may connect the two active sites (*23, 24, 26*).

**Figure 2:**
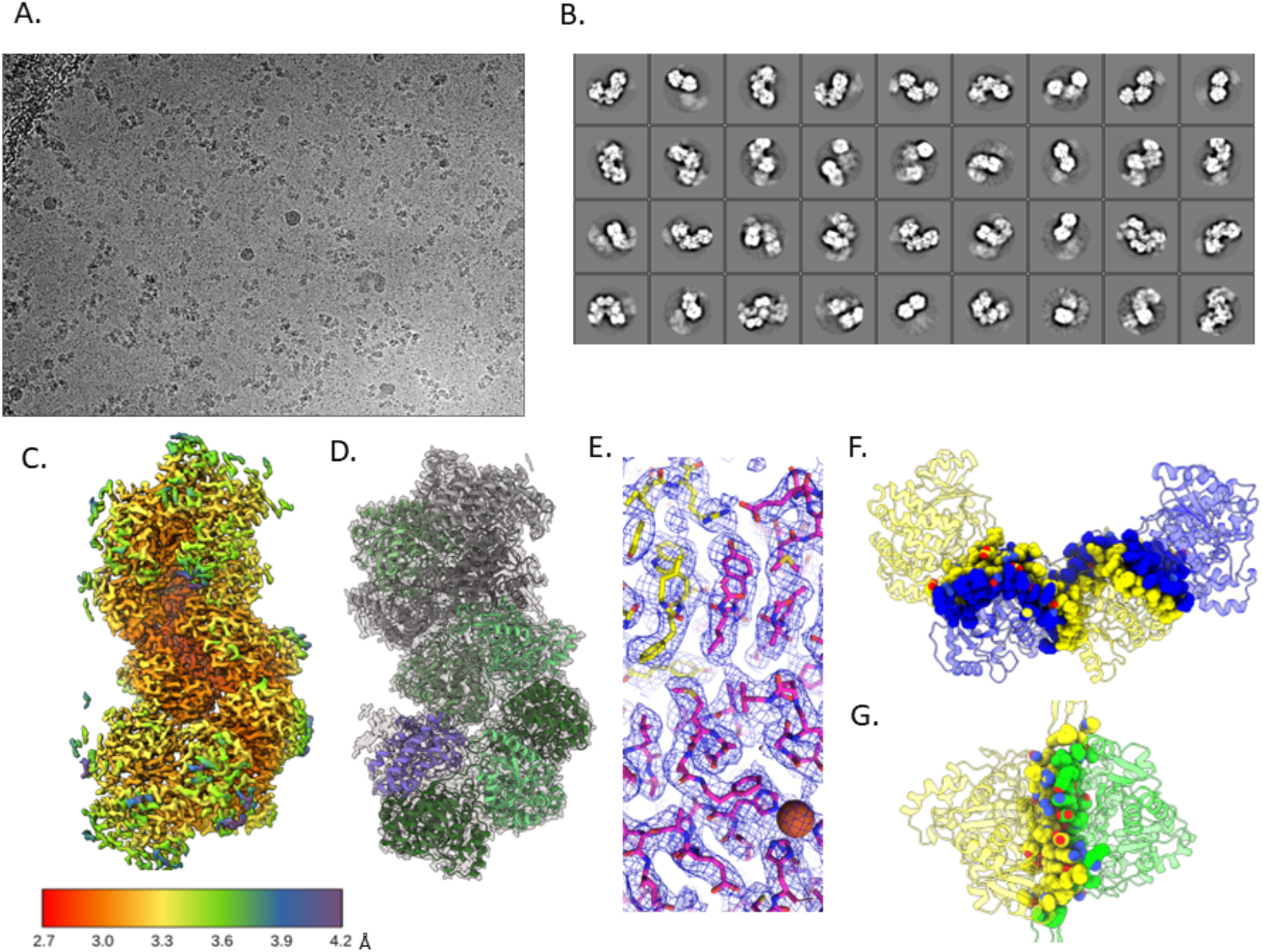
Cryo-EM structure of the extended *C. thermocellum* AdhE spirosome. **A)** Representative cryo-EM image **B)** 2D class averages of the single-particle analysis **C)** Local resolution of the final map **D)** Cartoon model fit in the density **E)** Stick representation to show that side chains can be determined at this resolution, the sphere represents the catalytic Fe in the ADH domain **F)** Dimer interface shows a swapped domain dimerization with a large buried surface area **G)** Tetramer interface has a smaller buried surface area.

### Structural comparison of spirosome oligomerization interfaces

There are two interfaces that form the spirosome (Figure 2 F, G). The first is the buried area located between two molecules of AdhE, in a domain-swapped dimer interface, where one ALDH domain is sandwiched between the ADH and ALDH domains of the second protein. This area contains a wide array of residues involved in the interaction that spans the entire protein (Figure 2F). The second interface is between the dimer of dimers, here referred to as the tetramer interface. This interface has a much smaller interaction area because it only occurs between two ADH domains (Figure 2G). Intriguingly, the overall hydrophobicity and electrostatics of these interfaces are similar between the *C. thermocellum* extended, *E. coli* extended, and *E. coli* compact structures (Supplemental Figure 2). However, the calculated buried surface area differs between the two *E. coli* dimer interfaces, but not the tetramer interface. Both the *E. coli* and *C. thermocellum* extended dimer interfaces bury ~5000 Å^2^. While the *C. thermocellum* compact dimer interface buries a similar surface area of ~4800 Å^2^, the *E. coli* compact dimer interface buries ~3800 Å^2^. Conversely, all the tetramer interfaces bury ~2000 Å^2^. One would expect the compact structure in *E. coli* to have a larger buried surface area due to it being the predominant form when it is examined without additives, but that is not the case; further corroborating that factors other than buried surface area must impact the *apo* state of the spirosome. Analysis of the residues buried in these interfaces reveals that while many of the residues are identical in the *C. thermocellum* and *E. coli* extended structures, there are some differences in amino acid type distribution, although nothing that directly indicates control of conformer state (Supplemental Figure 3).

We also examined the salt bridge and hydrogen bond networks between AdhE monomers (Figure 3). We found that more salt bridges form in the extended *C. thermocellum* structure as compared to the extended *E. coli* structure, which suggests that this may be a factor in the stability of extended *C. thermocellum* structure that is not found in the *E. coli* structure. However, when looking between the *E. coli* extended and compact structures, there are two more salt bridges in the extended structure than in the compact structure, which again implies that there might be additional factors impacting the primary form of the spirosome (*23, 26*). When looking at the hydrogen bonds between molecules, the extended *C. thermocellum* spirosome has a wider variety of hydrogen bonds (44 unique pairs) than the extended *E. coli* spirosome (32 unique pairs), indicating a potentially more stable structure. However, as with the salt bridges, the compact *E. coli* spirosome has fewer hydrogen bonds (17 unique pairs) than the extended structure, which does not explain why the compact form is the primary structure isolated when purified without substrates.

**Figure 3:**
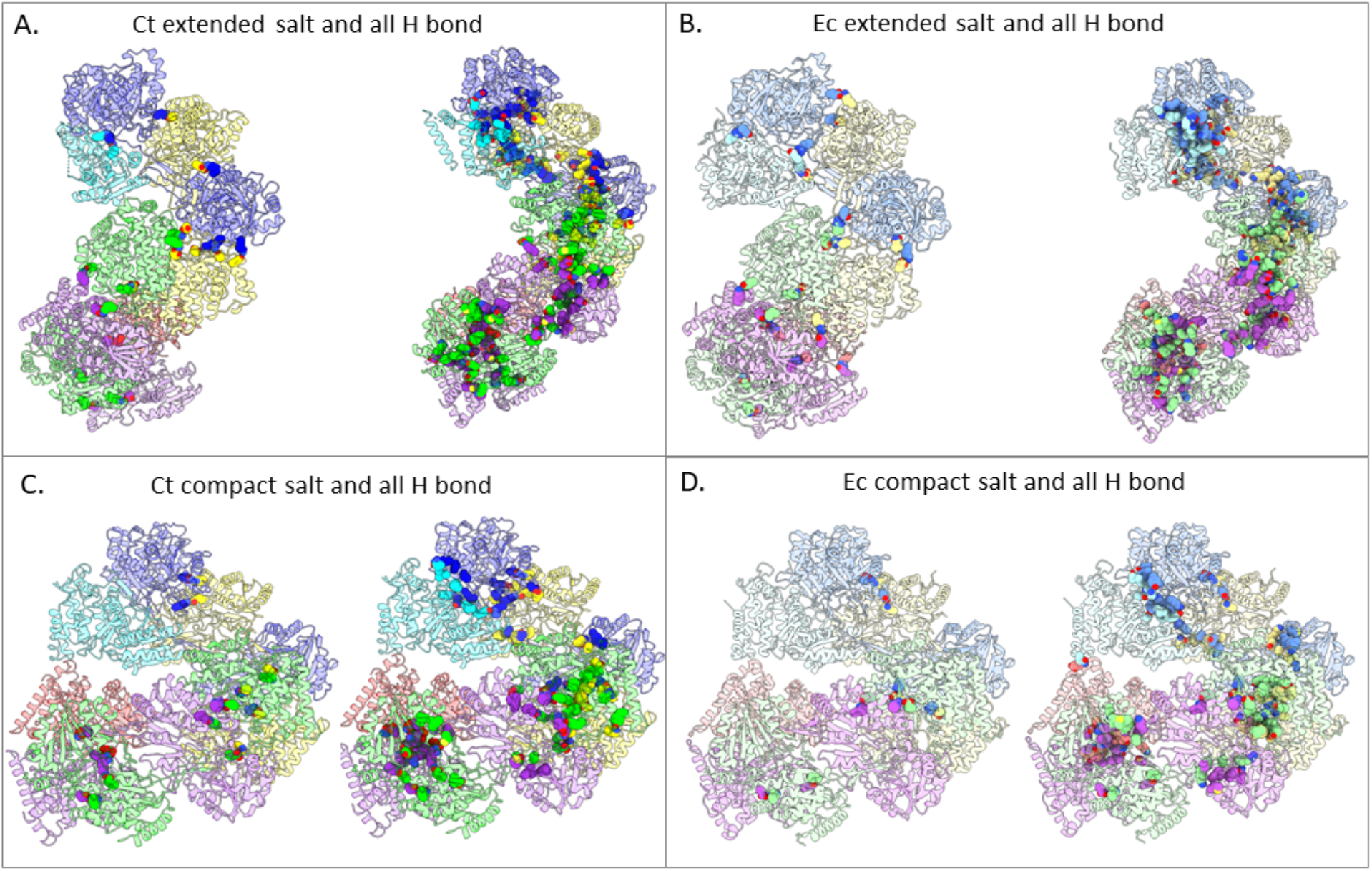
Hydrogen bond networks are more prevalent in the extended spirosomes of both *E. coli* and *C. thermocellum*. **A)** *C. thermocellum* extended spirosome. **B)** *E. coli* extended spirosome from PDB ID 7BVP. **C)** SWISS-MODEL of the *C. thermocellum* compact spirosome. **D)** *E. coli* compact spirosome from PDB ID 6AHC. Individual monomers have unique colors, the *E. coli* structure monomers are colored with lighter shades to correlate to the corresponding monomer in *C. thermocellum*. All salt bridges and hydrogen bonds are represented with spheres to emphasize their locations in the ribbon structure.

We also examined areas in the *C. thermocellum* structure that align with areas in the *E. coli* structures that have previously been shown to disrupt spirosome formation or with areas in the *V. cholerae* structure that were implicated in spirosome length. For the former in *E. coli*, there are two examples: 1) F670 mutated to either a small non-polar amino acid or a charged amino acid (PDBID 7BVP) and 2) the deletion of residues 446-449, both resulting in spirosome dissociation (PDBID 6TQH) (*23, 24*). In *C. thermocellum*, F670 is a tyrosine, however the hydrophobic pocket that surrounds that residue still exists (Figure 4A). We postulate that the similar shape of tyrosine maintains the integrity of this interaction, resulting in stable spirosomes. Regarding the 446-449 deletion, it occurs in the linker between the two domains of the AdhE monomer. This linker is vital for the domain-swapped structure of the dimer. Thus, deleting four residues, regardless of the identity of those amino acids, should interfere with the dimerization of AdhE (Figure 4B).

**Figure 4:**
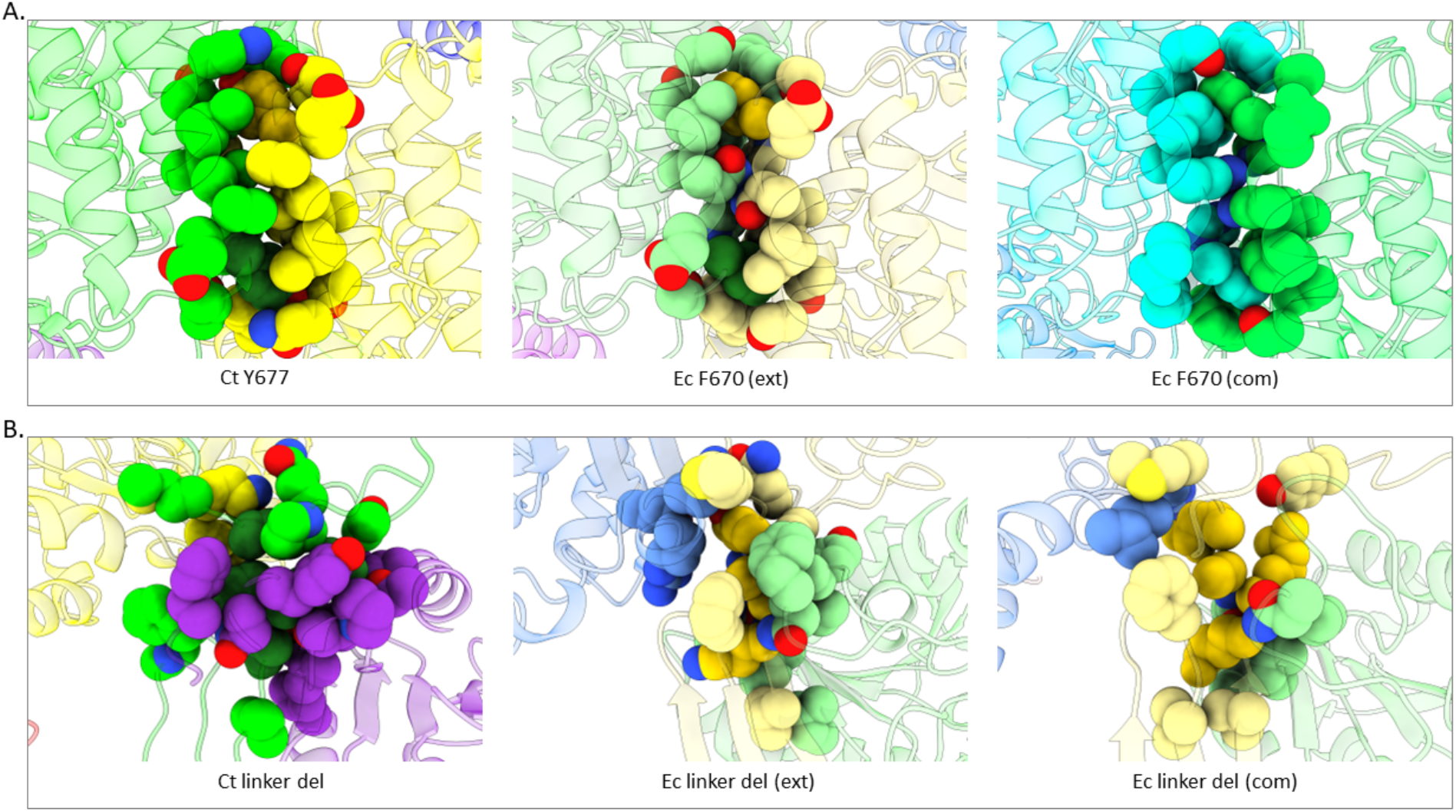
Previously identified mutants in *E. coli* compared to the *C. thermocellum* structure. **A)** Top row represents the mutant *E. coli* F670 that disrupts spirosomes, as well as *E. coli* F670/S705 that lengthens spirosomes as found in PDBID 7BVP. **B)** Bottom row represents the *E. coli* 446-449 linker deletion as found in PDBID 6TQH.

Finally, in *V. cholerae* AdhE expressed in *E. coli* (PDBID 7DAG), Cho *et al*. found that mutating Y669 and N704 to the corresponding residues in *E. coli*, phenylalanine and serine, resulted in longer spirosomes (i.e. more AdhE subunits in the spirosome) (*27*). The *C. thermocellum* sequence aligns with *V. cholerae*, with a tyrosine and asparagine in these positions (Figure 4A). However, based on our negative-stain results (Figure 1C), we show that *C. thermocellum* AdhE spirosomes, when expressed in *E. coli*, are significantly shorter than when in their native host. We extrapolate from these results that a similar phenomenon could occur if *V. cholerae* spirosomes were isolated from their native organism.

### Structural comparison of AdhE active sites

One factor that compaction and extension of the spirosome affects is the active site of both the ALDH and ADH domains. Here, we compare the NAD^+^-occupied *E. coli* site to the empty site of the *C. thermocellum* structure to determine which residues may interact with NAD^+^ (Figure 5A, B). In a sequence comparison between 25 AdhE proteins that have been previously identified in bacteria, both the ALDH and ADH NAD-pockets seem relatively well-conserved (Figure 5C) (*21, 22, 29-31*). There are a few residues with less than 50% identity (when using *C. thermocellum* as the basis of comparison), such as I119, L204, F496, S605, and M645. However, most of these residues retain similar chemistry at those locations - for example, L204 is a non-polar residue in every bacterium queried. Conversely, the amino acid that corresponds to the F496 position can vary between non-polar, negatively charged, and polar, which is surprising since the phenylalanine at that position stacks with the adenosine moiety of the NAD^+^. These patterns generally held true when the sequence alignment was expanded to 1000 AdhE sequences that were identified through the JGI IMG database (Supplemental Figure 4). Using this larger set of sequences, we also analyzed the D494 position. Prior work has shown that mutating the aspartate to glycine results in a change of co-factor preference from NADH to also include NADPH (*14*). There were 2.7% of AdhE sequences that had a naturally occurring G at this position, which suggests that there are AdhE proteins that rely on NADPH for their catalysis. We were able to visualize a potentially novel NAD(P)H binding site previously only identified via sequence comparison (Figure 5D) (*19*). The sequences contained a conserved GxGxxG motif at residues 426-431, which has been shown as involved in nucleotide binding (*32*). Intriguingly, this site does not coincide with either the ALD or ALDH active sites and is therefore less likely to have a catalytic function. However, this motif is located at the interface of two AdhE monomers, indicating that it could have an allosteric role, although further studies are needed to confirm. The three glycines at this position are all over 99% conserved, indicating that they play an important role in the AdhE structure or function.

**Figure 5:**
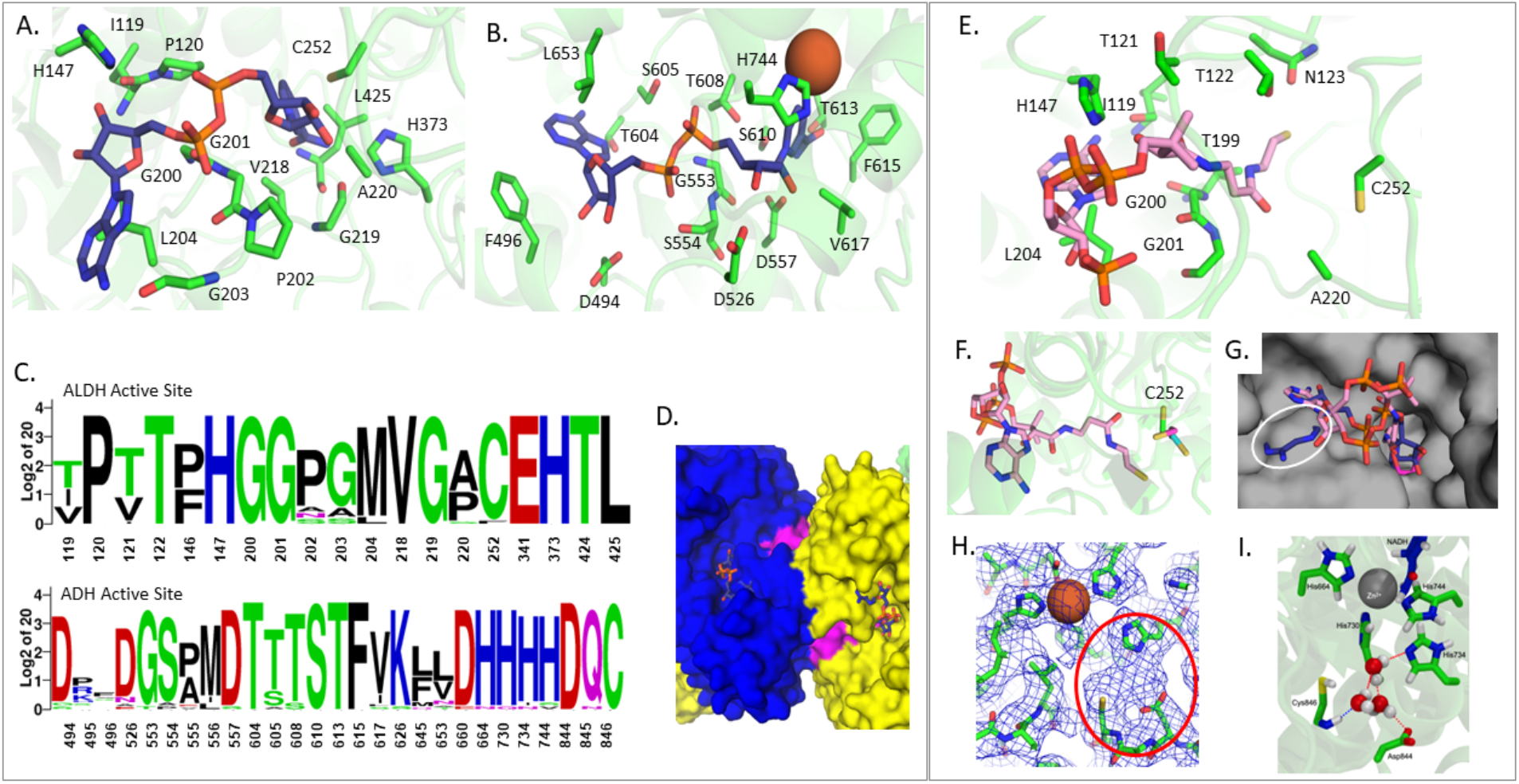
Active sites of *C. thermocellum* AdhE compared to *E. coli* AdhE. **A)** ALDH NAD^+^-binding domain, where all residues within 2.5 Å of the NAD^+^ are shown in stick (Ct - green, NAD^+^ - navy) **B)** ADH NAD^+^ binding domain, where all residues within 2.5 Å of the NAD^+^ are shown in stick (Fe^2+^ – rust sphere) **C)** Sequence alignment of the active site residues **D)** Surface representation of the novel NAD(P)H binding site (magenta) shown between two AdhE monomers (blue and yellow) **E)** ALDH binding domain compared to *R. palustris* (Rp) bound to acetyl-CoA, where all residues within 2.5 Å of the docked Rp acetyl-CoA are shown as sticks (Ct – green, acetyl-CoA – pink) **F)** Zoom panel from C showing the catalytic cysteine in relation to the docked acetyl-CoA (Ct -green, Ec – cyan, Rp – magenta) **G)** Surface view of *C. thermocellum* with both NAD^+^ (Ec, navy) and acetyl-CoA (Rp, pink) docked into the structure, white circle indicates a clash between NAD^+^ and *C. thermocellum* **H)** Density of the *C. thermocellum* ADH active site, showing the coordinated Fe^2+^ atom and an empty density between H734, D844, and C846, circled in red **I)** Snapshot from molecular dynamics simulation at the ADH active site, illustrating hydrogen bonding between water molecules occupying the active site and the three catalytic residues (H734, D844, and C846, green sticks). Also shown are NADH molecules (navy sticks), H644, H730, and H744 (green sticks), and zinc (gray sphere).

We also used structure alignments to estimate how both the *C. thermocellum* and *E. coli* ALDH active sites would accommodate an acetyl-CoA molecule *via* comparison to the structure of the *R. palustris* aldehyde dehydrogenase bound to acetyl-CoA (Figure 5E) (*33*). We observed that the proteins aligned with very little deviation - less than 2 Å RMSD – causing the acetyl-CoA aligned from *R. palustris* to sit in most of the same pocket that the NAD^+^ occupies as aligned from *E. coli*. This suggests that the dehydrogenase reaction occurs stepwise to include both the NAD^+^ and the acetyl-CoA (Figure 5G) (*33-35*). Furthermore, when looking at the surface representation of the *C. thermocellum* structure, we found that the NAD^+^ clashes with the protein, indicating that the pocket might shift to accommodate NADH separate to acetyl-CoA (Figure 5G). In looking at the residues within 3.5 Å of the acetyl-CoA, we identified multiple regions that seem conserved between all three organisms. Interestingly, if we look at the catalytic cysteine, which is covalently linked to the acetyl-CoA in the *R. palustris* structure, we see that the cysteine for both *C. thermocellum* and *E. coli* are rotated approximately 120° away from the *R. palustris* position in opposite directions which could be due to the absence of acetyl-CoA in these structures (Figure 5F) (*23, 33*).

At the ADH active site, we observe an empty density in the *C. thermocellum* extended structure, coordinated between residues that have been implicated in the ADH activity of the protein (Figure 5H, circled in red) (*16*). As illustrated in Figure 5F, we see that there is density, most probably a water molecule, being coordinated by H734, D844, and C846. D844 and C846 are 100% conserved, while H734 is 92% conserved amongst the 25 sequences that we analyzed. While slightly less well-conserved in the 1000 sequence comparison, all three residues are still highly conserved (H734 is 97.8%, D844 is 98.8%, and C846 is 91.6% conserved; Supplemental Figure 4). Furthermore, this small pocket is within 10 Å of the catalytic iron atom, which increases the probability that this region is important for catalysis. No published structure of the ADH domain contains either acetaldehyde or ethanol at the active site, though the structure of the *Geobacillus thermoglucosidasius* ADH domain (PDBID 3ZDR) contains a glycerol molecule coordinating a divalent metal ion (*29*). The terminal alcohol group of the glycerol molecule was speculated to be similar to that of ethanol, thus the structure may constitute a product mimic (*29*). Further supporting our hypothesis that a water molecule may occupy the observed empty density at the ADH active site in the *C. thermocellum* extended structure, two independent MD trajectories, each with two full AdhE molecules including two ADH actives sites, demonstrate hydrogen bonding of at least one water molecule with D844 in essentially every frame (>98%), as well as with H734 (16-31% of the frames) and with C846 (7-66% of frames with both side chain and main chain, as shown in Figure 5I.

### Identification of a channel that confines aldehydes and examination of the dynamics of spirosome ultrastructures

Intermediate channeling has been proposed previously in the *E. coli* spirosome, based on observation of enclosed space within the structure, and has been hypothesized as a functional role to contain reactive aldehydes and therefore limit cytotoxicity (*24*). However, direct evidence for the functional role of this putative channel is still lacking. In order to more fully understand the extended spirosome of *C. thermocellum*, we employed multiple computational approaches to analyze its structure. First, we used MOLE 2.0 to identify potential channels within the structure of the spirosome, specifically looking for channels that span between the ALDH and ADH active sites (*36*). In all three structures (*E. coli* compact, *E. coli* extended, and *C. thermocellum* extended), as well as a SwissPDB model of the *C. thermocellum* compact structure, MOLE 2.0 identified a channel that could connect the two active sites. Compelling trends emerge when analyzing the composition of amino acids that line the channel compared to the amino acid composition of the whole protein for both *E. coli* and *C. thermocellum*. Overall, the channels are more highly conserved than the overall protein: the residues lining the extended channel have 92% homology, and the residues in the compact channel have 97% homology, whereas the overall protein homology is only 62%. When more AdhE sequences are included in the analysis, more variety is seen in the residues lining the channel (Figure 6A, Supplemental Figure 5). Further examination of the individual amino acids also reveals important trends. First, there is only one lysine in the tunnel in all three structures, showing one of the largest log2-fold changes as compared to the composition of the full protein (Figure 6B). This may indicate that the channel is designed to contain reactive aldehydes while avoiding cross-linking with the amine group of the lysine. Second, there is a higher concentration of histidines in the channel in all the structures as compared to the full protein. This is potentially due to the tunnel passing by the catalytic iron in the ALDH domain, which is coordinated by three histidines. Notably, there is a difference in the number of phenylalanines in the channel when compared between the compact and extended structures. The extended structures contain a much higher density of phenylalanines as indicated by the log2-fold change. Finally, we focused on the cysteines present in these channels. In the compact state, only the catalytic cysteine from the ALDH domain is present in the channel. However, in both extended structures, there are two cysteines in the channel: the catalytic cysteine from the ALDH domain and the cysteine in the catalytic pocket from the ADH domain. This could point to the extended spirosome as being the relevant catalytic form because there is a potential path that efficiently connects the active sites.

**Figure 6:**
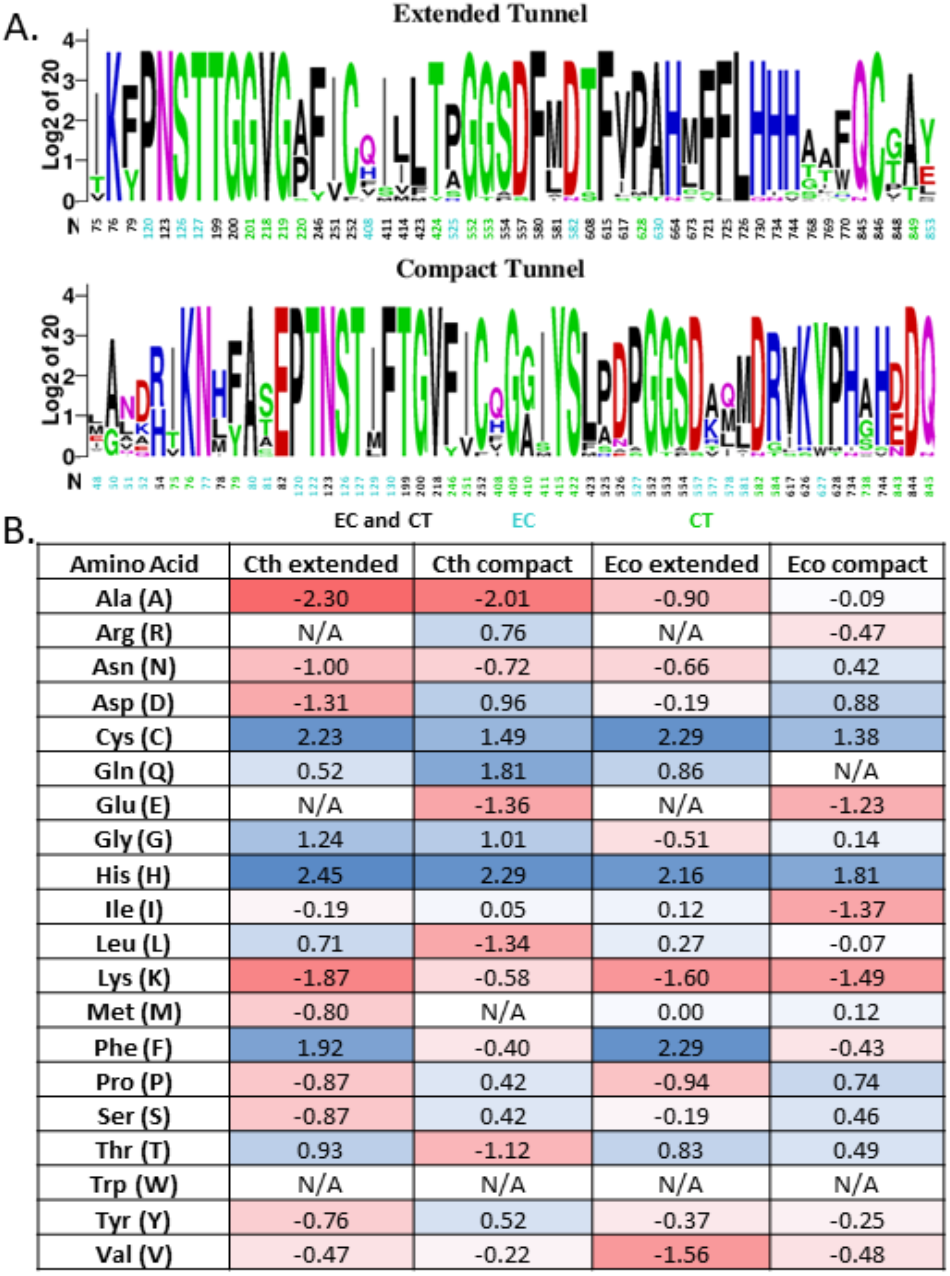
Analysis of the residues lining the spirosome channel. **A)** Sequence homology of the channel from both *C. thermocellum* and *E. coli*, residues only identified in the Ct channel are green and in the Ec only channel are cyan. **B)** Log_2_-fold difference between the residues that line the channel and the full-length protein. The scale of change is color-coded from deep red for fewer residues represented in the channel to deep blue for more residues represented in the channel.

As a complement to the insights gained from the static structures, molecular dynamics (MD) simulations with bound acetaldehyde were conducted to investigate the dynamics of the aldehyde intermediate. We examined the effect of spirosome configuration (extended and compact) and species (*C. thermocellum* and *E. coli*) on residence time of acetaldehyde in the enzyme before escape to the solvent. The starting enzyme configurations for these simulations were the cryo-EM structure of the extended *C. thermocellum* spirosome, a homology model of the *C. thermocellum* compact spirosome (PDBID 6TQM serves as the template), and *E. coli* spirosomes in extended (6TQH) and compact (6TQM) forms (*24*). As in 6TQH and 6TQM, all simulation systems comprise two full AdhE units and two additional ADH domains on either side. The aldehyde intermediate is formed at the ALDH active site, thus this constitutes the starting location for acetaldehyde in the simulations. Additional starting points midway between the ALDH and ADH catalytic sites were also generated based on the results from MOLE, namely midway along the MOLE tunnel where the channel was the widest (Figure 7, panels A and C). More details for the system construction, acetaldehyde starting points, and simulation details are found in the SI.

**Figure 7:**
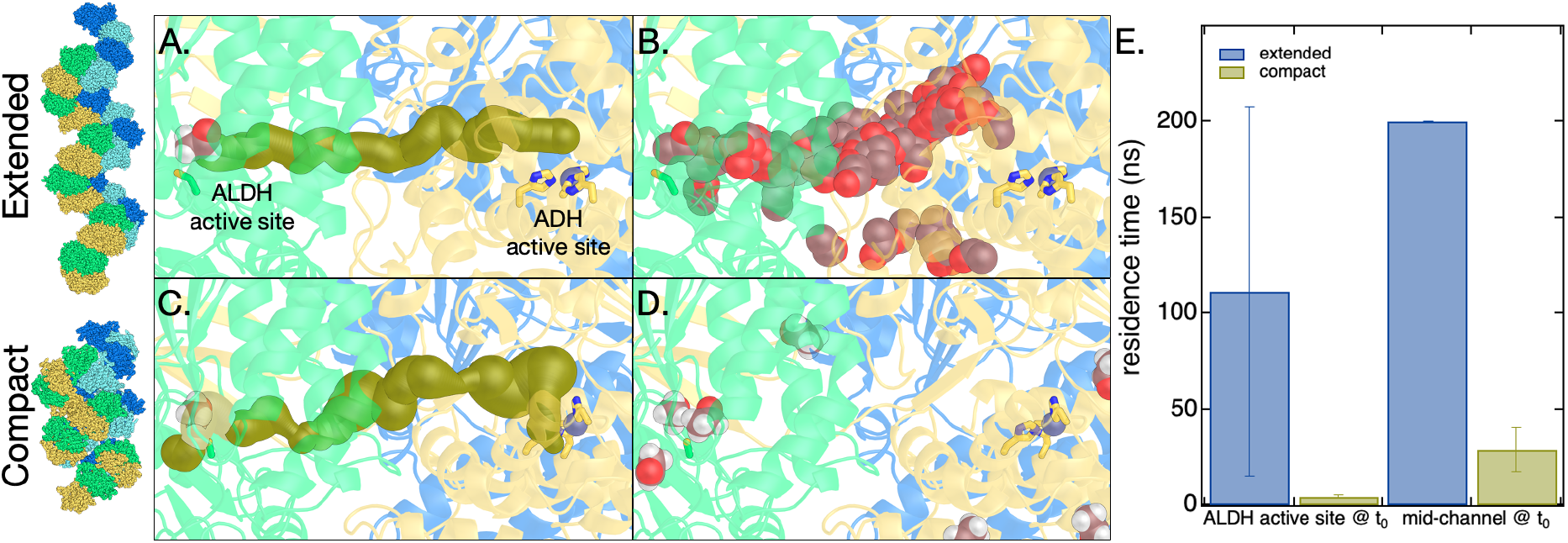
Molecular dynamics (MD) of aldehyde channeling in C. thermoceullum AdhE spirosomes. **A)** Starting configuration for MD simulation of the *C. thermocellum* extended spirosome structure overlaid with the channel connecting the ALDH and ADH active sites as determined by MOLE (shown in gold spheres) (36). Also shown are C252 (representing the ALDH active site, green sticks), three histidine residues that coordinate the divalent metal at the ADH active site (His664, His730, His744, yellow sticks), zinc ion (purple sphere), the starting location for acetaldehyde (violet spheres), and the tertiary AdhE structure (each AdhE molecule colored distinctly). **B)** Same as panel A, except MOLE tunnel is removed, and acetaldehyde location from MD simulation is shown every 1 ns for 160 ns. In this simulation, the acetaldehyde molecule exits the enzyme into solution after about 125 ns. **C)** Representations are the same as in A, but here the channel shown is determined by MOLE for the compact spirosome structure (*E. coli* 6TQM and aligned with *C. thermocellum* AdhE homology structure). **D)** Same as panel C, except MOLE tunnel is replaced with acetaldehyde location from MD simulation, shown every 1 ns. In this simulation, the acetaldehyde molecule exits the enzyme into solution at about 4 ns. **E)** Average residence time for acetaldehyde in channel before exiting AdhE in MD simulations for two different starting configurations for the AdhE spirosome (extended and compact) and acetaldehyde (at ALDH active site and midway along the channel). Error bars are the standard deviation for simulations performed in triplicate.

Three 200 ns MD simulations were conducted from each starting configuration. The primary conclusion from these is that the residence time for acetaldehyde when the spirosome is in the extended conformation is significantly longer than the compact conformation (Figure 7E, Supplemental Figure 6E). The significant difference between extended and compact forms presented in Figure 7E could be even larger given that many of the 200 ns simulations ended with acetaldehyde still present within the extended AdhE spirosomes (and thus the true residence time is longer than 200 ns). In addition, the trajectory of the aldehyde tracks quite closely with the channel found by MOLE (Figure 7A, 7B). Though the acetaldehyde is sometimes observed to escape AdhE when started in the extended conformation, it typically samples a significant stretch of the path between the ALDH and ADH active sites, occasionally traversing the full distance between the two active sites (Supplementary Movie 1). In contrast, simulations initiated with the spirosome in the compact conformation typically see acetaldehyde leave within 4 ns (ALDH active site starting position) to 30 ns (starting point midway in the channel). For example, in the trajectory represented in Figure 7D acetaldehyde leaves within 4 ns. We conducted analogous simulations with the *E. coli* AdhE and found the basic conclusions to be analogous to the *C. thermocellum* AdhE (Supplementary Figure 6).

Taken together, these simulations effectively present a novel and dynamic understanding of intermediate aldehyde channeling in AdhE spirosomes and lend evidence to the hypothesis that the extended spirosome may be the dominant active complex for AdhE spirosomes.

Finally, we turned back to the variability of spirosome conformation observed by negative-stain TEM and searched more closely for the same variability in the cryo-EM data. Initial visual analysis was unable to discern between 2D projections of extended or compact conformations, as the difference was expected to be minute. The abundance of extended spirosomes relative to compact spirosomes confounded computational cross-correlation sorting while simultaneously making it difficult to visually identify compact projections. To address this, we sorted all particles selected by the crYOLO autopicker (which exhibits less bias than template or manual picking) using cryoDRGN (*37*).

Correlation between particles visualized graphically in the UMAP (Figure 8A) indicated the existence of at least four distinct spirosome conformations. Volumes generated from these particle subgroups indicated the presence of extended, compact, and at least two major intermediate conformations (Figure 8B). We attempted to refine the isolated classes from cryo-DRGN but encountered extended spirosome contamination in each of the classes. However, we continued searching for compact spirosome projections by generating 2D templates from the *E. coli*, compact AdhE spirosome model (PDBID 6AHC) using cryoSPARC (Figure 8C) (*23, 38*). These templates were then used as a visual reference, which did indeed reveal the presence of compact 2D projections within the *C. thermocellum* cryoEM data (6.7% of picked particles were compact) that were previously overlooked (Figure 8D). The subsequent reconstruction using these particles yielded a 3.93 Å density of the compact AdhE spirosome, although there was evidence of streaking due to preferred orientation issues and low particle numbers. Although the streakiness prevented the successful fitting of a high-resolution molecular model, the density map corroborates the domain orientation of a compact spirosome, as modeled by SWISS-MODEL. There are some intriguing insights into differences between our compact ultrastructure compared to previously reported compact AdhE spirosomes isolated from different species (*23, 24, 27*). Most notably, the *C. thermocellum* AdhE does not achieve as compact a state, even when isolating the most compact particles. Moreover, the reconstruction of a compact spirosome density corroborates that compact spirosomes observed *via* negative staining are also present in near-native conditions (cryogenically preserved samples). This data confirms that spirosomes have the potential to exist in a spectrum of states that can be captured in a single cryo-EM grid without any additional cofactors to induce conformational changes.

**Figure 8:**
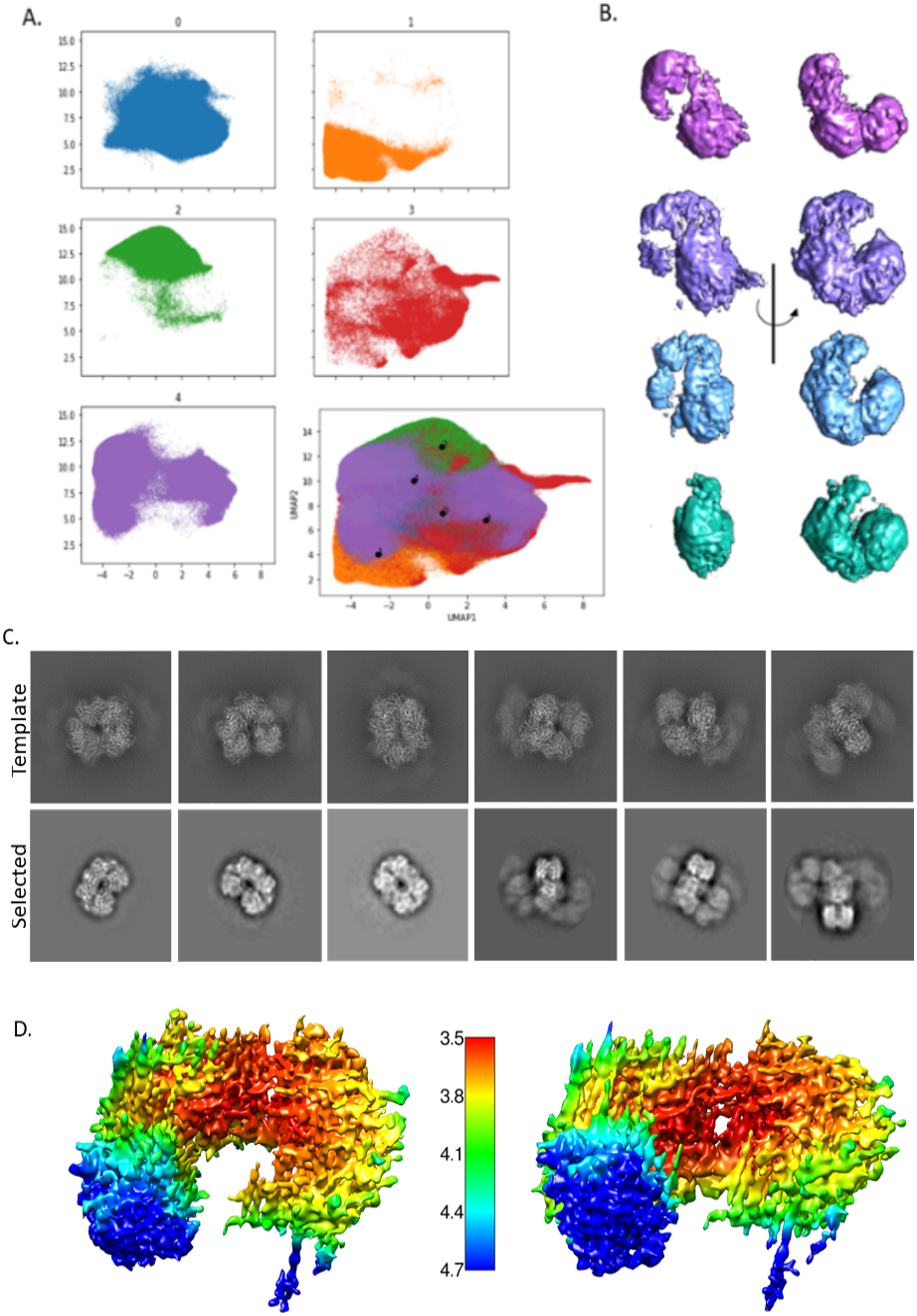
CryoDRGN results show that there is conformational heterogeneity in the sample. **A)** UMAP representation of the particles shows a continuous heterogeneity (bottom right corner). Colored UMAP graph shows the division of the particles into five classes, broken into their individual map based on color. **B)** One representative density for each group (excepting the junk class) shown at two angles to illustrate the movement of the spirosome (purple – extended, lavender – intermediate 1, light blue – intermediate 2, teal – compact). **C)** Back-projected template classes generated in Cryosparc from PDB ID 6AHC (top) and the matching classes selected from our data (bottom). **D)** Two angles of the final compact spirosome density shown with local resolution coloring as indicated by the scale bar in the center.

## Conclusions

In this study, we presented and analyzed a high-resolution structure of the AdhE spirosome from *C. thermocellum*. In comparison to the *apo E. coli* spirosome structure, which purifies in the compact conformation, the *apo C. thermocellum* spirosomes are primarily extended (*23, 24*). Our structural comparison identifies the extended conformation as the more structurally stable form in both the case of *E. coli* and *C. thermocellum*, which disagrees with the reality of the compact spirosome being the stable, *apo* conformation from *E. coli*. The evidence for the extended spirosome as the more stable form includes the higher number of salt bridges and hydrogen bonds between molecules in the extended as compared to the compact. Furthermore, extended spirosomes have a larger buried surface area in the dimer interface than the compact spirosomes (5000 Å^2^ compared to 3800 Å^2^, respectively), imparting additional stability. Although we could not structurally identify exact features that would predict which conformation was preferred, we have evidence from negative-stain EM experiments that conformation does not rely solely upon the environment: *C. thermocellum* spirosomes are extended in both endogenous and exogenous expression conditions. Therefore, there must be some intrinsic structural elements that contribute to the *apo* state of the spirosome, although we have not found that element in this study. This could potentially be due to the differences between Gram-positive and Gram-negative bacteria. In previous studies, compact spirosomes have only been isolated from Gram-negatives while solely extended spirosomes have been isolated from Gram-positives. Furthermore, while the compact spirosomes can transition to extended in the presence of cofactors, the reverse has not been previously observed with an extended spirosome (*22-24, 31*). Here, we identified compact spirosomes in our study, although they were largely outnumbered by the more-preferred extended spirosome, which further indicates that the extended structure is more stable for *C. thermocellum*, regardless of expression host. While these compact spirosomes could result from expression in *E. coli*, which is very unlikely, we also identified compact spirosomes in a native *C. thermocellum* lysate, which would not have similar contamination issues. Close examination of the structure revealed a potential novel allosteric NAD(P)H binding site that lies between two monomers of AdhE, identified by a characteristic NAD(P)H binding motif. The GxGxxG motif is highly conserved across 1000 AdhE sequences, indicating its importance in the protein. Further studies will be necessary to fully understand the role of this motif.

The second conclusion of this paper is that intermediate channeling is possible and may be the primary purpose of large, spirosome ultrastructures. An intriguing finding from prior research is that the spirosome is necessary for the acetyl-CoA to acetaldehyde reaction in the ALDH domain, yet unneeded for the reverse reaction or for ADH activity (*23, 24*). Therefore, a channel might be necessary to funnel acetaldehyde away from the ALDH active site to prevent product inhibition in the ALDH site. Prior research identified a similar channel in *E. coli* AdhE spirosomes yet did not confirm its role nor characterize the movement of intermediates within the channel (*24*). Here, we identified an enclosed tunnel between an ALDH domain and an ADH domain of two separate proteins in *C. thermocellum* and used MD to examine possible aldehyde channeling. While the channel can be found in both the compact and extended structures, we see that the channel in the compact conformation has areas that are exposed to the cytosol, whereas the extended channel is entirely enclosed. This further supports the idea that the extended spirosome is the active form of AdhE (*26*). MD simulations further corroborate this, showing that the aldehyde intermediate has a much longer occupation time in the channel of the extended form. Future mutagenesis studies will be needed to confirm whether the spirosome exists to control the reaction flux in high-reactant conditions.

The *C. thermocellum* AdhE structure, in combination with molecular dynamics and comparative analysis with the *E. coli* structures, establishes a baseline for further studies using mutagenesis to increase the efficacy of the protein. This work provides insight as to why the spirosome seems to be necessary for only one direction of the reaction, as well as opens new and interesting questions as to why certain variants of AdhE natively exist in different conformations. It also provides a template that could be replicated in other systems where toxic intermediates need to be sequestered to increase the production of valuable biochemicals.

## Materials and Methods

### Gene cloning in E. coli

*E. coli* and *C. thermocellum* AdhE genes fused with N-terminal His-tag and TEV site used in this study were amplified from their genomic DNA and inserted into pET28b vector. Gibson Assembly Cloning Kit (NEB, Ipswich, MA, USA) and 5-alpha *E. coli* strain (NEB, Ipswich, MA, USA) were used for the gene cloning. Gene cloning protocols reported previously were followed (*39*). Primers, plasmids, and strains used in the study are listed in Supplemental Table 1. (To examine the protein sequences of the expressed genes, please reference Supplemental Figure 7).

### Gene expression in E. coli

The generated plasmids used for AdhE gene expression described above were transformed into *E. coli* BL21(DE3) strain (NEB, Ipswich, MA, USA). The transformants were grown aerobically in LB medium supplemented with kanamycin at 37 °C and 225 rpm overnight (seed culture). The seed culture in a 1:50 ratio was inoculated into LB medium containing the same antibiotic and the cells were grown under the same growth conditions until optical cell density OD_600_ reached 0.8. IPTG (0.3mM) was added to the culture, and cells were grown overnight at 16 °C and 225 rpm after the culture was cooled on ice for 3 hours. The cells were collected by centrifuge 8000xg for 10 min and used for further protein extraction and purification.

### Protein Isolation and Purification

Cell lysis was performed in HisA buffer (50 mM Tris pH 7.5, 400 mM NaCl, 20 mM imidazole, 1% glycerol), 20 mg of lysozyme, 1 uL of DNaseI, and one protease inhibitor tablet, all set to rock at room temperature for one hour. Lysed cells were spun down at 15,000× g, and the clarified lysate was loaded onto a His-trap column for affinity purification. The tagged protein was eluted in HisB (50 mM Tris pH 7.5, 400 mM NaCl, 400 mM imidazole, 1% glycerol), then applied to an S200 size exclusion column for a final step of purification in 50 mM Tris pH 7.5 and 150 mM NaCl. Protein was stored at 4°C before either negative stain or cryo-EM grid preparation.

### Native C. thermocellum cell growth and isolation

Fermentations were carried out using *Clostridium thermocellum* strain DSM1313. Cultures were grown in Medium for Thermophilic Clostridia (MTC), using stock solutions to combine with water and the desired carbon source to reach 1x concentration. The medium was prepared as described with cellobiose as the carbon source (*40*).

Glycerol stocks were revived in 10 mL serum bottles that were prepared in an anaerobic chamber containing 85% N_2_, 10% CO_2_, and 5% H_2_. Seeds were grown to 0.8-1.0 OD_600_ at 55°C and transferred to 100 mL serum bottles. These cultures were grown to an OD_600_ of 0.8-1.0 in approximately 12 hours before the entire contents (10% v/v) were inoculated into 5 g/L cellobiose MTC medium in process control bioreactors for a final working volume of 1 L. All growth medium was prepared at pH 7.0 and maintained at this level for all bioreactor experiments using 2N KOH. Temperature was maintained at 55°C with 50 rpm agitation for bottles and bioreactors. An N_2_ environment was maintained through the bioreactor headspace at a rate of 30 mL/min.

Cultures were grown until maximum cell density was reached around 1.2 OD_600_ in approximately 8 hours. The entire bioreactor contents were harvested into centrifuge bottles and centrifuged at 8,000 RPM for 20 minutes. The supernatant was removed by decanting. The pellet was then washed with distilled water and centrifuged again under the same conditions. The wash fraction was removed by decanting, and the residual pellet was frozen until use in experiments.

Cells were lysed using a BeadBeater with a 30 seconds on/30 seconds off pattern for 5 minutes in SEC buffer (50 mM HEPES pH 7.4, 150 mM NaCl) with 1 uL of DNaseI and one tablet of protease inhibitor. Lysate was spun down at 15,000× g to pellet any extra cell debris, then concentrated, and passed through a 0.22 um filter before being loaded directly onto a Superose6 size exclusion column. The fractions that had signal on the FPLC trace were stored at 4C before negative stain grid preparation.

### Negative-Stain TEM Sample Preparation and Data Collection

All AdhE proteins were screened for optimal concentrations for negative staining – finding that 0.05 mg/ml was the ideal concentration for protein separation. Grids were glow discharged for 20 seconds at 10 mA before the sample was applied to the grid to incubate for one minute. The excess sample was blotted away before the grid was dipped in a droplet of 2% aqueous uranyl acetate (UA) for 15 seconds. Excess UA was blotted away, and grids were left to dry before storage at room temperature before transmission electron microscopy (TEM) screening. Grids of the *apo* nCthAdhE, CthAdhE, and EcoAdhE were viewed on a Tecnai T20 (FEI/ThermoFisher Scientific) TEM at 50,000x magnification, with 6x6 montages collected using SerialEM (*41*). Grids of the CthAdhE and EcoAdhE with added reactants were viewed using a Talos L120C TEM at 36,000x magnification, using an automated collection strategy through Leginon (*42*).

### Negative Staining Data Analysis

Spirosome length and overall conformation were manually analyzed in Image J (FIJI) (*43*). Image pixel size was used to calibrate the scale bar before the line tool was used to trace and measure the length of the spirosome. Compact versus extended was determined by eye for at least 100 and up to 1200 spirosomes for statistical relevance. Statistical relevance of differences in length of spirosomes was calculated using a Mann-Whitney U Test, all were found to be statistically different from each other.

### Cryo-EM Sample Preparation and Data Collection

C-Flat 1.2/1.3 400 mesh grids were glow discharged for 10 seconds before 0.4 – 0.5 mg/mL of CthAdhE was applied. Grids were blotted for 3 seconds using a CP3 Cryoplunge 3 (Gatan Ametek Inc.) before being plunged into liquid ethane held at -168 °C. Preliminary data was collected on a 200kV Talos Arctica (ThermoFisher Scientific) cryo-TEM with K3 direct electron detector (Gatan Ametek Inc.) using a total electron dose of 90 e^-^/Å^2^ and an exposure time of 3 seconds. The data set used in the final extended *C. thermocellum* spirosome reconstruction was collected on a 300 kV Titan Krios cryo-TEM with a Falcon 4 direct electron detector (ThermoFisher Scientific) using a total electron dose of 60 e^-^/Å^2^ and an exposure time of 6.49 seconds. Finally, the data used for the compact CthAdhE analysis was collected on a 300 kV Titan Krios cryo-TEM with a Falcon 4 (ThermoFisher Scientific) direct electron detector using a total electron dose of 60 e^-^/Å^2^ and an exposure time of 4.42 seconds.

### Cryo-EM Data Analysis

Structural analysis was done in Relion 3.1, originally starting with helical processing to create a 3D template for autopicking in single particle analysis, resulting in a 3.8 Å structure that was used for initial model building (Supplemental Table 2, Supplemental Figures 8 and 9) (*44*). Data from the Krios was also processed in Relion 3.1 using single particle analysis (*44*). Using a soft mask around the edge of the protein, a final map was generated that had an FSC 0.143 of 3.28 Å. The CCP-EM software suite was used to dock a homology model of the *C. thermocellum* structure into the density (*45*). Coot was used to refine the structure iteratively with CCP-EM refinement and validation (*46*). The best structure was taken to Phenix, where it was docked into the auto-sharpened map generated by Phenix (*47*). Final rounds of refinement and validation resulted in the structure deposited to the PDB as 8UHW and the final map was deposited to the EMDB as EMD-44284.

### Sequence Alignments and Analysis

25 sequences previously identified in literature were aligned in Clustal Omega (*21, 22, 29-31, 48*). The percentage of conservation was calculated by hand and the alignment logos were generated with WebLogo (*49*). For the 1000-sequence dataset, sequences were identified by homology on the JGI IMG database (*50*). All relevant sequences were input into the MAFFT online service (*51, 52*). Protein length settings were used to exclude sequences that only aligned with one domain before final analysis of the consensus sequences.

### Molecular Dynamics Simulations

Classical MD simulations of 200 ns in length were run in triplicate for the following enzymes: 1) *C. thermocellum* AdhE in extended spirosome form, 2) *C. thermocellum* AdhE in compact form, 3) *E. coli* AdhE in extended form, and 4) *E. coli* AdhE in compact form. With the exception of *C. thermocellum* in compact form, the starting structures for the enzymes come from cryo-EM structures. The *E. coli* extended and compact forms originate with the structures from Pony *et al*. (PDB codes 6TQH and 6TQM, respectively) (*24*). The *C. thermocellum* compact spirosome was constructed as a homology model by SWISS-MODEL with 6TQM as the template (*24, 28*). The simulation system comprises two full AdhE units and two additional ADH domains on either side, as in 6TQH and 6TQM. Where they appear in the cryo-EM structures, NAD+ and NADH molecules are removed. All three cryo-EM structures contain divalent iron bound at ADH active sites. The position of these cations is maintained, but iron is swapped for zinc in each case. We have chosen to include zinc in each ADH active site, rather than iron, for two primary reasons. The first is that parameters for iron are not included in the standard CHARMM forcefield. Secondly, it has been shown that ADH activity of ADHE can be stimulated in cell extracts to varying degrees, including zinc (*29*). In fact, divalent zinc has been shown to have higher ADH activity than divalent iron in at least one study, leading the authors to conclude that this class of ADH structures should be described as “metal ion-dependent” rather than strictly iron- or zinc-specific (*29*).

In the biofuels context, i.e. producing alcohols from acyl-CoA molecules, the aldehyde intermediate is formed at the ALDH active site from acyl-CoA and then travels to the ADH active site where it is reduced to an alcohol. Thus, for our MD simulations, positioning the acetaldehyde intermediate at the ALDH active site most closely approximates the physical situation for intermediate channeling. The placement of acetaldehyde at the ALDH active site was determined by alignment with 5JFM (*33*). Additional starting points midway between the ALDH and ADH catalytic sites were generated based on the results from MOLE, namely choosing starting coordinates midway along the MOLE tunnel where the channel was widest (Supplemental Figure 5F, G) (*36*).

In general, protonation states were determined by the output from H++ server (http://biophysics.cs.vt.edu/H++) at pH 6.0, consistent with the assays performed by Extance *et al*. In addition, the catalytic cysteine residues in ALDH domains (C252 in *C. thermocellum and* C246 in *E. coli*) are deprotonated. The overall charge on the enzymes domains (two, full AdhE molecules and two additional ADH domains) with these protonation states is -42 for *C. thermocellum* systems and -28 for *E. coli* systems; a corresponding number of sodium ions were added to the solution phase to neutralize the overall system. Neither *C. thermocellum* nor *E. coli* AdhE contains any disulfide bonds.

Furthermore, to investigate the dynamics of water molecules at the ADH active site, 80 ns MD simulations were performed in duplicate of the *C. thermocellum* extended structure with NADH bound at the ADH active site. Zinc molecules are positioned at each ADH active site, as above.

All simulations were built using CHARMM version 44a1 and simulated with the CHARMM36 force field for the protein, and TIP3P potential for water molecules (*53-55*). Topologies and forcefield parameters for acetaldehyde and NADH come from the CHARMM generalized forcefield (CGenFF) version 4.1 (*56, 57*).

### CryoDRGN Analysis

To evaluate structural heterogeneity evident in the cryoEM data, 959,561 potential spirosome particles were sorted into 2D classes using RELION 3.1 and were exported for analysis with cryoDRGN 0.3.4 (*37, 44*). Particles were downsampled to a 64px box size [5.28 Å/px] and analyzed for 50 epochs with the standard latent space parameter of 8, at which point training converged. Inspection of 20 volumes generated by k-means clustering and resulting UMAP (Uniform Manifold Approximate and Projection) visualization of latent space of particle projections suggested the presence of four major conformational classes, which were segmented by Gaussian mixture modeling (GMM, k=4). Of them, only major classes indicative of spirosome formation were retained for further analysis. A fifth class (‘Other’) of unidentifiable “junk” particles and dimers was also identified, and those particles were discarded from further processing. The remaining 858,511 spirosome particles were scaled to 128px box size [2.64 Å/px] and again analyzed for 50 epochs. Major classes indicated by the UMAP clusters were attributed to extended, compact, and transitional state spirosomes by visual confirmation of 3D density volumes generated by k-means clustering and filtered by GMM (k=5).

### CryoSPARC Template Generation and Use

The cryoSPARC ‘Create Templates’ utility was used to back project 50 2D classes of the compact *E. coli* AdhE spirosome (*24, 38*). For posterity, 50 back projected 2D classes of the extended *E. coli* AdhE spirosome were also generated to serve as a comparison (*24*). Visual inspection indicated sufficient difference between 2D projections of both conformations. The 2D templates of the compact structure were used as a visual reference by which compact classes were hand selected from the heterogenous compact model building described above (Supplemental Table 1). This resulted in the selection of 64,977 particles that were determined to be in the compact state, which were processed in Relion 3.1 as described above. Because the heterogenous dataset was lacking in some projections predicted by the template generation, the resulting 3D density of the compact *C. thermocellum* AdhE suffered from some preferred orientation and ultimately achieved a 3.93 Å resolution, as determined by the Relion gold standard FSC.

## Acknowledgments

Portions of this work were performed at the CU Anschutz Medical Campus Cryo-EM Facility and the SLAC S2C2 Cyro-EM Facility. We thank both facilities for the use of their microscopes and related equipment. Computer time was provided by the National Renewable Energy Laboratory Computational Sciences Center supported by the DOE Office of EERE under contract number DE-AC36-08GO28308.

## Author Contributions

Conceptualization: SJZ, BCK, YJB

Methodology: SJZ, NH, QX, EER, LMJ, BCK

Investigation: SJZ, BCK, JNG

Visualization: SJZ, BCK, JNG

Supervision: YJB

Writing—original draft: SJZ, BCK, JNG

Writing—review & editing: SJZ, BCK, JNG, NH, QX, EER, LMJ, YJB

## Data and materials availability

The molecular model of AdhE can be found at PDB ID 8UHW. The refined map, half-maps, and FSC curves can be accessed at the EMDB EMD-42284. All other data is accessible in the main text or supplementary information.

