## Supplemental Information for "Structural characterization and dynamics of AdhE ultrastructures from *Clostridium thermocellum:* A containment strategy for toxic intermediates"

Samantha J. Ziegler *et al.*

**This PDF file includes:**

Figs. S1 to S9
Tables S1 and S2
Movie S1

**Other Supplementary Materials for this manuscript include the following:**
Movie S1

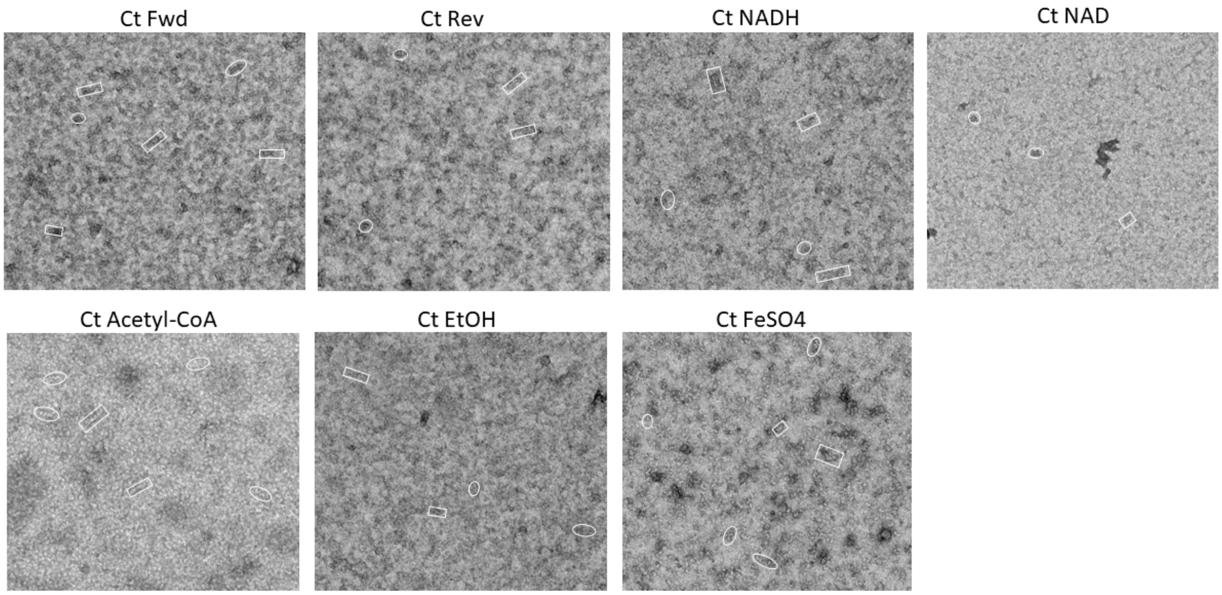


**Fig. S1. Negative stain images of C. thermocellum AdhE in the presence of reactants**

A representative image from each of the added reactants from Figure 1B. Each image has compact spirosomes in rectangles and extended spirosomes in ovals. Fwd – Acetyl-CoA, NADH, FeSO4; Rev – Ethanol, NAD+, FeSO4.

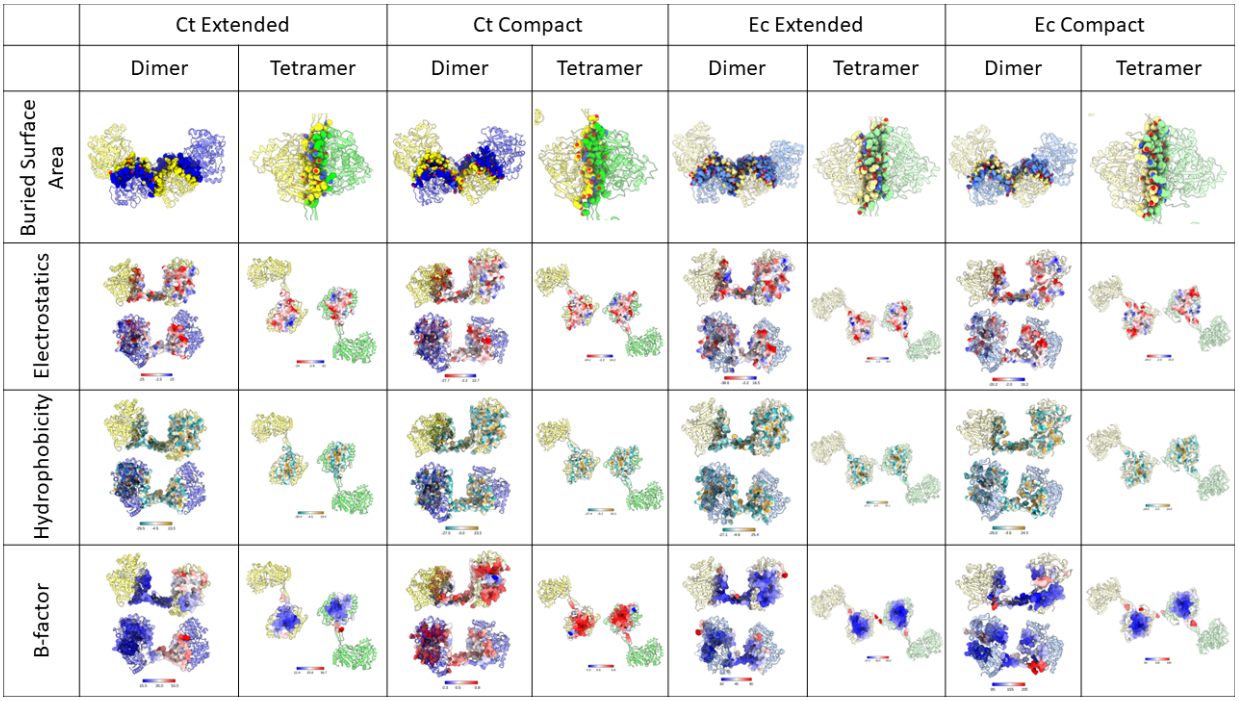


**Fig. S2. Comparing structural properties of the interfaces of the *C. thermocellum* and *E. coli* AdhE spirosomes**

A chart examining the dimer and tetramer interfaces of the *C. thermocellum* (Ct) and *E. coli* (Ec) extended and compact structures. Ec extended – PDBID 7BVP, Ec compact – PDBID 6AHC, Ct compact – SWISS Model. Scale bars for the electrostatics, hydrophobicity, and B-factor are included at the bottom of each cell.

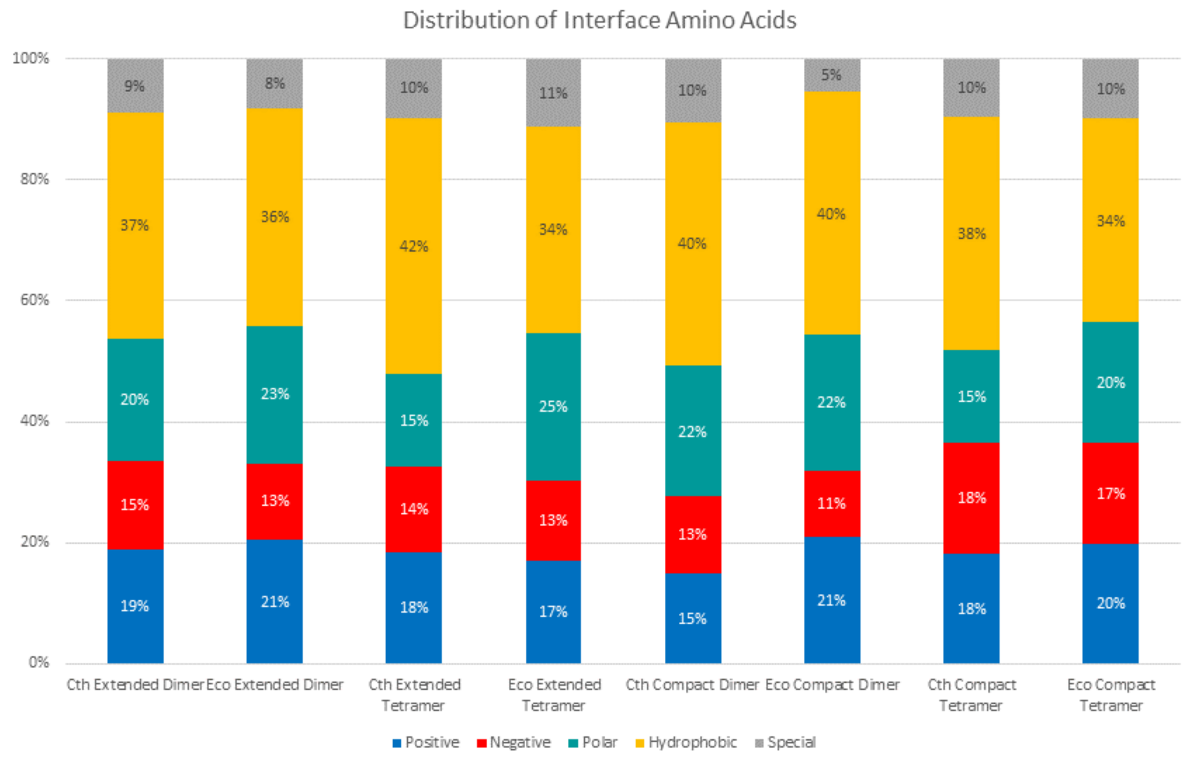


**Fig. S3. Distribution of amino acids found in spirosome interfaces**

Bar graph representing the amino acid types found in the interfaces. “Special” amino acids are cysteine, glycine, and proline.

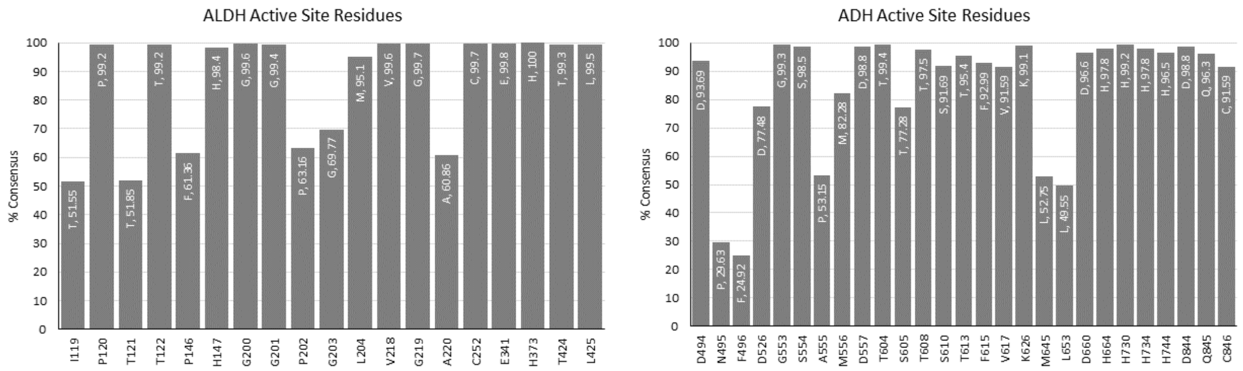


**Fig. S4. AdhE active site consensus sequences**

Graphs showing the prevalence of residues at positions lining the NADH binding pocket of both the ALDH and ADH domains in 1000 AdhE sequences. The position on the x-axis represents the location in *C. thermocellum*. The residue listed at the top of each bar indicates the consensus residue, as well as the percentage of consensus.

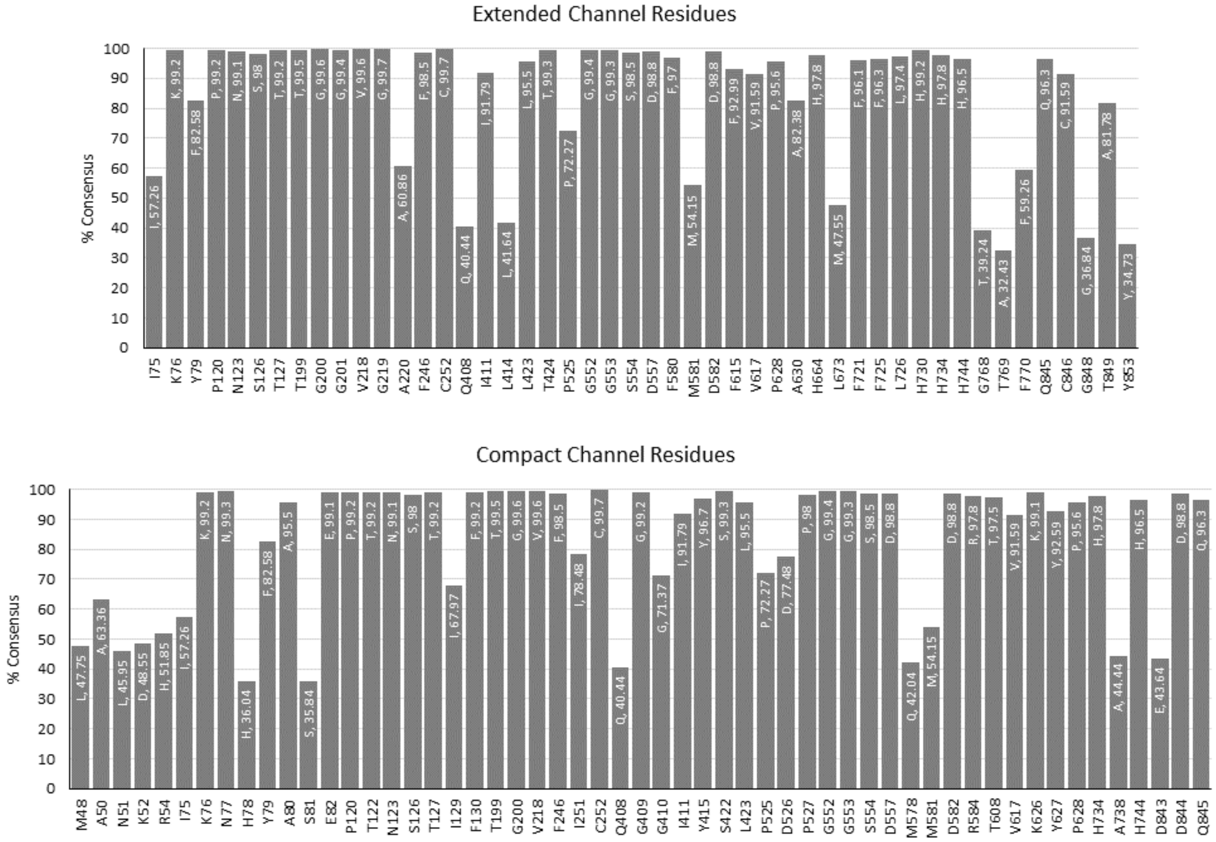


**Fig. S5. Consensus sequences of the AdhE channel-lining residues**

Graphs showing the prevalence of residues at positions lining the extended and compact channels in 1000 AdhE sequences. The position on the x-axis represents the location in *C. thermocellum*. The residue listed at the top of each bar indicates the consensus residue, as well as the percentage of consensus.

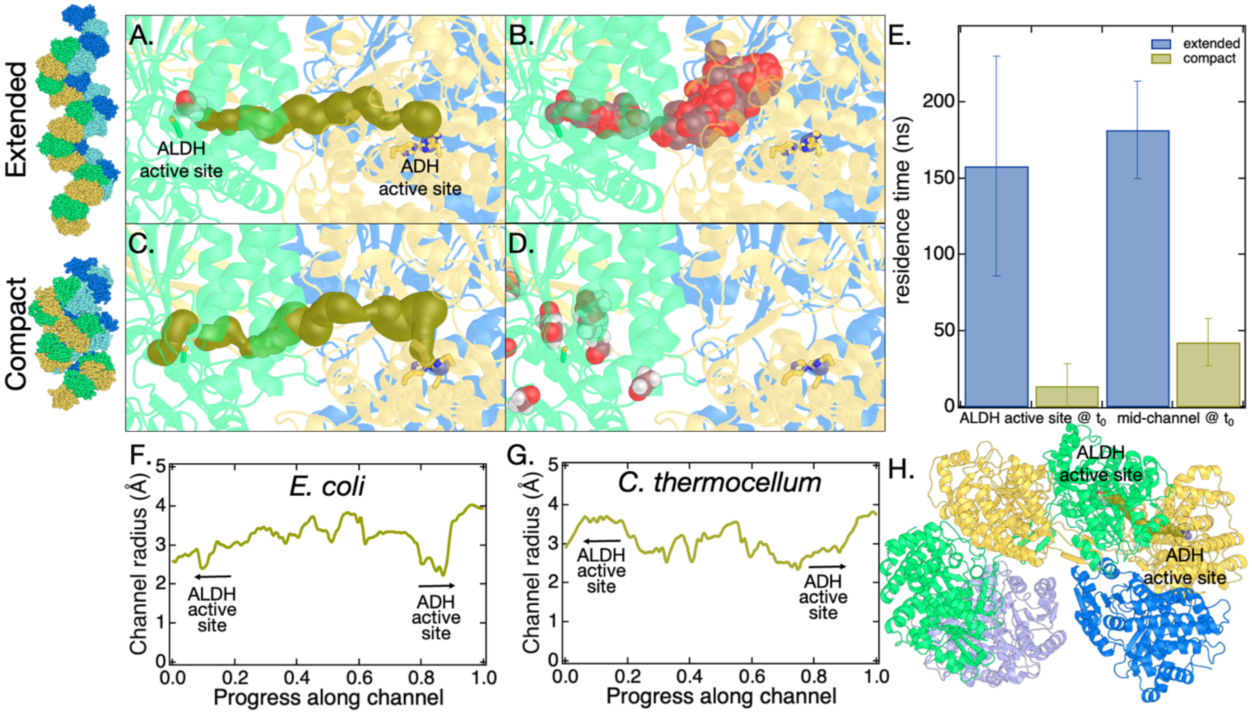


**Fig. S6. Molecular dynamics (MD) of aldehyde channeling in *E. coli***
**A)** Starting configuration for MD simulation of the *E. coli* extended spirosome structure overlaid with the channel connecting the ALDH and ADH active sites as determined by MOLE *(36)* (shown in gold spheres, structure used is PDB code 7BVP). Also shown are C246 (representing the ALDH active site, green sticks), three histidine residues that coordinate the divalent metal at the ADH active site (His657, His723, His737, yellow sticks), zinc ion (purple sphere), the starting location for acetaldehyde (violet spheres), and the tertiary AdhE structure (each AdhE molecular colored distinctly). **B)** Same as panel A, except MOLE tunnel is removed, and acetaldehyde location from MD simulation is shown every 1 ns for 200 ns. In this simulation, the acetaldehyde molecule remains within the enzyme for the full 200 ns simulation. **C)** Representations are the same as in A, but here the channel shown is determined by MOLE for the compact spirosome structure (structure used is PDB code 6TQM and aligned with homology structure for compact *C. thermocellum* spirosome). **D)** Same as panel C, except MOLE tunnel is replaced with acetaldehyde location from MD simulation, shown every 1 ns. In this simulation, the acetaldehyde molecule exits the enzyme into solution at about 8 ns. **E)** Average residence time for acetaldehyde in channel before exiting AdhE in MD simulations for two different starting configurations for the AdhE spirosome (extended and compact) and acetaldehyde (at ALDH active site and midway along the channel). Error bars are the standard deviation for simulations performed in triplicate. **F)** Channel radius profile from MOLE for *E. coli* extended cryo-EM structure (PDB code 6TQH). **G)** Channel radius profile from MOLE for *C. thermocellum* extended cryo-EM structure. **H)** To give a sense for the overall MD system setup, the extended *E. coli* spirosome (PDB code 6TQH) is shown as an example (solvent omitted for clarity). Two full AdhE molecules are shown in yellow and green. Light blue and royal blue indicate the two capping ADH domains. The active site for ALDH and ADH domains are indicated. Also shown is the tunnel found by MOLE that connects the two active sites. This panel is analogous to panel A, shown at larger scale.

### >E. coli AdhE in pCB16-1

```
MHHHHHHENLYFQGMAVTNVAELNALVERVKKAQREYASFTQEQVDKIFRAAALAAADARIPLAKMAVAESGMGIVEDKVIKNHFASEYIYNAYK
DEKTCGVLSEDDTFGTITIAEPIGIICGIVPTTNPTSTAIFKSLISLKTRNAIIFSPHPRAKDATNKAADIVLQAAIAAGAPKDLIGWIDQPSVE
LSNALMHHPDINLILATGGPGMVKAAYSSGKPAIGVGAGNTPVVIDETADIKRAVASVLMSKTFDNGVICASEQSVVVVDSVYDAVRERFATHGG
YLLQGKELKAVQDVILKNGALNAAIVGQPAYKIAELAGFSVPENTKILIGEVTVVDESEPFAHEKLSPTLAMYRAKDFEDAVEKAEKLVAMGGIG
HTSCLYTDQDNQPARVSYFGQKMKTARILINTPASQGGIGDLYNFKLAPSLTLGCGSWGGNSISENVGPKHLINKKTVAKRAENMLWHKLPKSIY
FRRGSLPIALDEVITDGHKRALIVTDRFLFNNGYADQITSVLKAAGVETEVFFEVEADPTLSIVRKGAELANSFKPDVIIALGGGSPMDAAKIMW
VMYEHPETHFEELALRFMDIRKRIYKFPKMGVKAKMIAVTTTSGTGSEVTPFAVVTDDATGQKYPLADYALTPDMAIVDANLVMDMPKSLCAFGG
LDAVTHAMEAYVSVLASEFSDGQALQALKLLKEYLPASYHEGSKNPVARERVHSAATIAGIAFANAFLGVCHSMAHKLGSQFHIPHGLANALLIC
NVIRYNANDNPTKQTAFSQYDRPQARRRYAEIADHLGLSAPGDRTAAKIEKLLAWLETLKAELGIPKSIREAGVQEADFLANVDKLSEDAFDDQC
TGANPRYPLISELKQILLDTYYGRDYVEGETAAKKEAAPAKAEKKAKKSA-
```

### >C. thermocellum AdhE in pCB17-12

```
MHHHHHHENLYFQGMTKIANKYEVIDNVEKLEKALKRLREAQSVYATYTQEQVDKIFFEAAMAANKMRIPLAKMAVEETGMGVVEDKVIKNHYAS
EYIYNAYKNTKTCGVIEEDPAFGIKKIAEPLGVIAAVIPTTNPTSTAIFKTLIALKTRNAIIISPHPRAKNSTIEAAKIVLEAAVKAGAPEGIIG
WIDVPSLELTNLVMREADVILATGGPGLVKAAYSSGKPAIGVGAGNTPAIIDDSADIVLAVNSIIHSKTFDNGMICASEQSVIVLDGVYKEVKKE
FEKRGCYFLNEDETEKVRKTIIINGALNAKIVGQKAHTIANLAGFEVPETTKILIGEVTSVDISEEFAHEKLCPVLAMYRAKDFDDALDKAERLV
ADGGFGHTSSLYIDTVTQKEKLQKFSERMKTCRILVNTPSSQGGIGDLYNFKLAPSLTLGCGSWGGNSVSDNVGVKHLLNIKTVAERRENMLWFR
TPEKIYIKRGCLPVALDELKNVMGKKKAFIVTDNFLYNNGYTKPITDKLDEMGIVHKTFFDVSPDPSLASAKAGAAEMLAFQPDTIIAVGGGSAM
DAAKIMWVMYEHPEVDFMDMAMRFMDIRKRVYTFPKMGQKAYFIAIPTSAGTGSEVTPFAVITDEKTGIKYPLADYELLPDMAIVDADMMMNAPK
GLTAASGIDALTHALEAYVSMLATDYTDSLALRAIKMIFEYLPRAYENGASDPVAREKMANAATIAGMAFANAFLGVCHSMAHKLGAFYHLPHGV
ANALMINEVIRFNSSEAPTKMGTFPQYDHPRTLERYAEIADYIGLKGKNNEEKVENLIKAIDELKEKVGIRKTIKDYDIDEKEFLDRLDEMVEQA
FDDQCTGTNPRYPLMNEIRQMYLNAYYGGAKK-
```

**Fig. S7. Sequences of proteins expressed in this manuscript**
The full sequences of the *E. coli* and *C. thermocellum* AdhE genes. The 6-His tag is highlighted in cyan and the TEV protease cleavage site is highlighted in green.

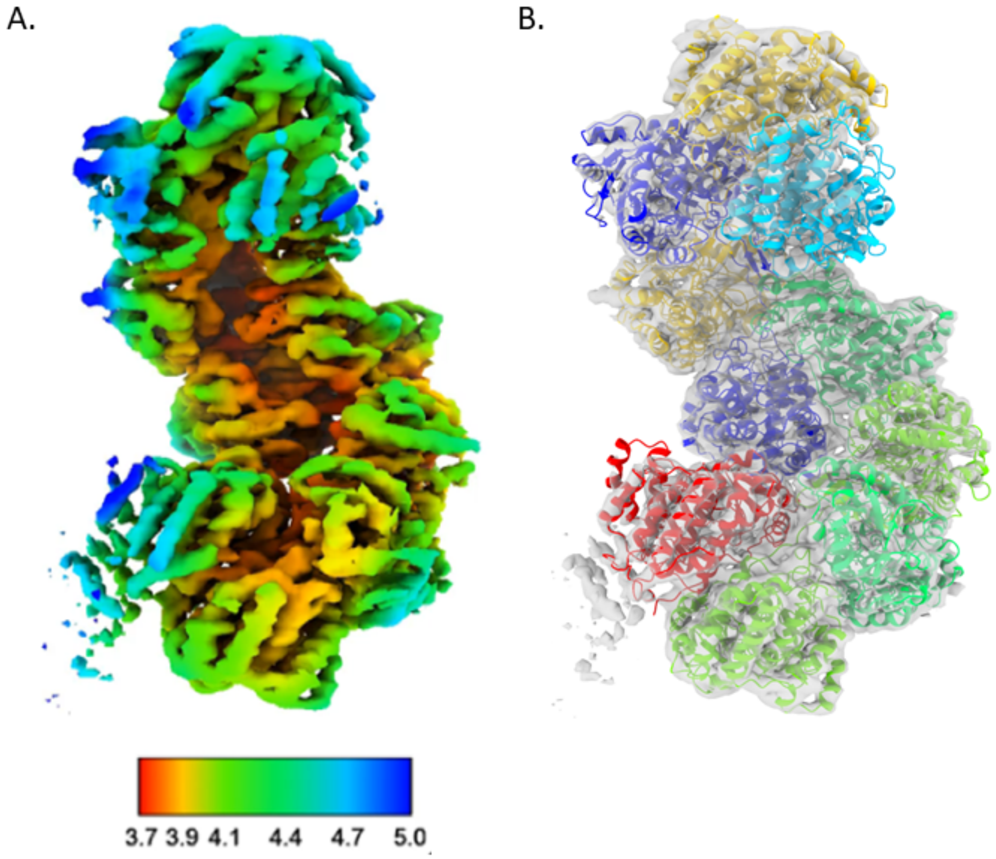


**Fig. S8. Lower-resolution *C. thermocellum* AdhE structure**
**A)** Density of the 3.8 Å AdhE structure colored by local resolution. **B)** Initial model generated fit into the density, gray. This was used as the template for the final structure model.

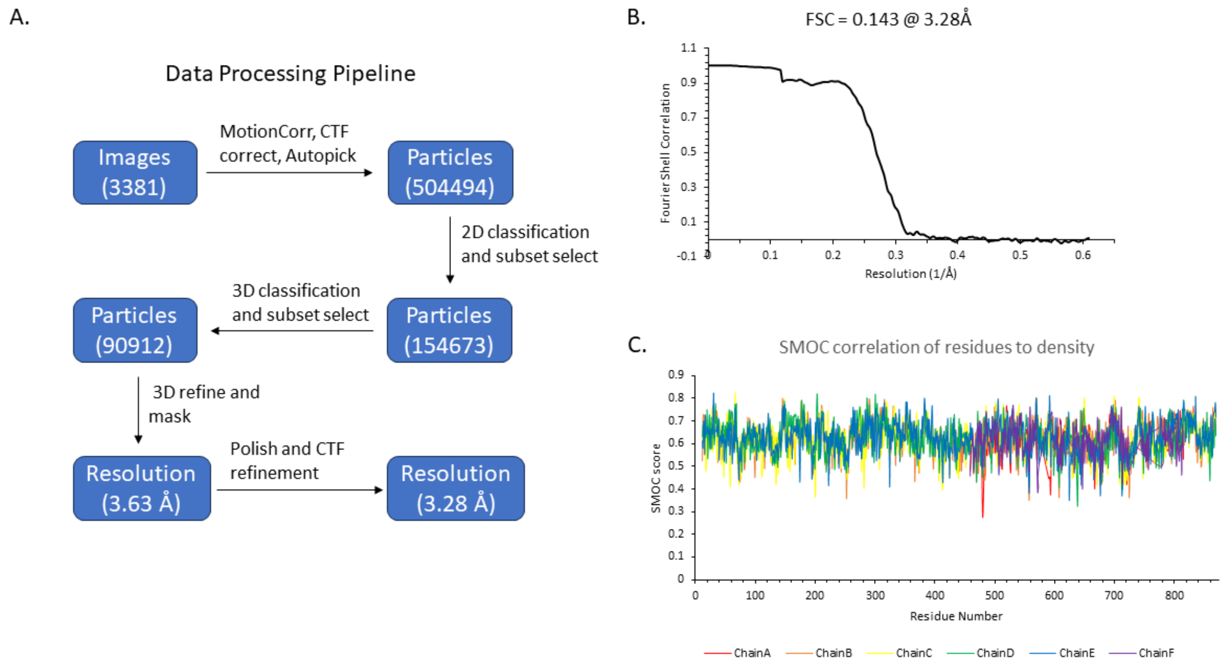


**Fig. S9(*1-4*). Data processing of the *C. thermocellum* AdhE structure**

**A)** Pipeline of data processing indicating number of particles at each major step, as well as resolutions pre- and post-polishing. **B)** Final FSC curve of the masked density, indicating the structure is 3.28 Å at and FSC of 0.143. **C)** SMOC correlation of the residue to density fit for all 6 chains as calculated by TEMPy local in CCPEM.

| Strain | Plasmid | Target gene | Genetic tag | Primers used for cloning | |
|---|---|---|---|---|---|
| | | | | Name | Sequence |
| | | | | 1184 | tgagatccggctgctaacaaa |
| | | | | 1252 | ccggatatagttcctcctttcag |
| | | | | 1524 | CATGGTATATCTCCTTCTTAAAGTTAAACAA |
| | | | | 1710 | taattttgtttaactttaagaaggagatataccatgcaccaccaccaccaccacgagaacctgtatttccagggtatgGCTGTTACTAATGTCGCTGA |
| | | | | 1711 | ctcagcttcctttcgggctttgttagcagccggatctcaAGCGGATTTTTTCGCTTTTTTCTC |
| | | | | 1712 | CGTTCCGACCACTAACCCGAC |
| | | | | 1713 | GCTGAACTGGCAGGCTTCTCT |
| M7-2 | pCB16-1 | E. coli AdhE | N-histag | 1714 | TTCGAAGTAGAAGCGGACCCG |
| | | | | 1184 | tgagatccggctgctaacaaa |
| | | | | 1252 | ccggatatagttcctcctttcag |
| | | | | 1524 | CATGGTATATCTCCTTCTTAAAGTTAAACAA |
| | | | | 1715 | ttttgtttaactttaagaaggagatataccatgcaccaccaccaccaccacgagaacctgtatttccagggtatgACGAAAATAGCGAATAAATACGAAG |
| | | | | 1716 | ctcagcttcctttcgggctttgttagcagccggatctcaTTTCTTCGCACCTCCGTAATA |
| | | | | 1717 | GTTGTGCCCGGTACTGGCAAT |
| | | | | 1571 | TTATAAGCCCACACCCCAGGG |
| | | | | 1572 | GTAGCTGACGGTGGATTTGGCC |
| M8-2 | pCB17-12 | C. thermocellum AdhE | N-histag | 1573 | GACACCATAATTGCGGTCGGC |

**Table S1. Plasmids, primers, and strains used in this study**

| | Extended Final | Extended Model Building | Compact Model Building |
|---|---|---|---|
| **Data collection** | | | |
| Microscope/camera | Krios/Falcon 4 | Talos Arctica/K3 | Krios/Falcon 4 |
| Voltage (kV) | 300 | 200 | 300 |
| Magnification | 96,000 | 45,000 | 96,000 |
| Electron dose ($e^-/Å^2$) | 60 | 90 | 60 |
| Pixel Size (Å) | 0.82 | 0.89 | 0.82 |
| Defocus Range (µm) | 1.5-3.0 | 1.2-2.5 | 1.0-3.0 |
| Processing | Single particle analysis | Single particle analysis | Single particle analysis |
| Symmetry imposed | C1 | C1 | C1 |
| Initial particle images (no.) | 504,494 | 352,061 | 959,561 |
| Final particle images (no.) | 90,912 | 103,973 | 64,977 |
| Map resolution (Å)-(FSC threshold model) | 3.28-(0.143) | 3.8-(0.143) | 3.93-(0.143) |
| **Refinement and validation** | | | |
| Map sharpening (B-factor) ($Å^{-2}$) | -97.02 | -99.00 | |
| **Model composition** | | | |
| No. of chains | 11 | 10 | |
| Atmos (no.) | 30,785 | 31,285 | |
| Residues (no.) | 3,994 | 4,047 | |
| Ligands (no.) | $Fe^{2+}$ (5) | $Fe^{2+}$ (4) | |
| Bond lengths (Å) | 0.003 | 0.004 | |
| Bond angles (˚) | 0.590 | 0.633 | |
| Ramachandran favored % | 95.26 | 93.92 | |
| Ramachandran allowed % | 4.74 | 6.01 | |
| Ramachandran outliers % | 0.00 | 0.08 | |
| Rotamers outliers % | 0.00 | 0.00 | |
| MolProbity score | 1.65 | 1.92 | |
| Clashscore | 5.84 | 9.80 | |
| CC (mask) | 0.86 | 0.79 | |
| CC (box) | 0.70 | 0.70 | |
| CC (peaks) | 0.70 | 0.62 | |
| CC (volume) | 0.82 | 0.78 | |
| Mean CC for ligands | 0.84 | 0.77 | |

**Table S2. Data collection and processing**

**Movie S1. Molecular dynamics (MD) of aldehyde channeling.**

Example of MD simulation of *C. thermocellum* extended spirosome with the channel connecting the ALDH and ADH active sites as determined by MOLE *(36)* shown in gold spheres. Also shown are C252 (representing the ALDH active site, green sticks), three histidine residues that coordinate the divalent metal at the ADH active site (His664, His730, His744, yellow sticks), zinc ion at ADH active site (purple sphere), and the tertiary AdhE structure (each AdhE molecular colored distinctly). Acetaldehyde (violet spheres) is initiated at the ALDH active site and throughout the simulation for 130 ns. In this simulation, the acetaldehyde molecule exits the enzyme into solution after about 125 ns. The movie was created in Pymol (The PyMOL Molecular Graphics System, Version 2.0 Schrödinger, LLC) and the positions of each residue are smooth using the "smooth" command with two passes and a window size of two. This same starting configuration was run in triplicate, with retention times of 9 and 200 ns in the other two simulations.


1. G. Kim *et al.*, Aldehyde-alcohol dehydrogenase forms a high-order spirosome architecture critical for its activity. *Nat Commun* **10**, 4527 (2019).
2. R. Laurenceau *et al.*, Conserved Streptococcus pneumoniae spirosomes suggest a single type of transformation pilus in competence. *PLoS Pathog* **11**, e1004835 (2015).
3. S. O. Matayoshi, H., Detection of Fine Spiral Structures (Spirosomes) by Weak Sonication in Some Bacterial Strains. *Microbiol Immunol* **29**, 13-20 (1985).
4. P. Pony, C. Rapisarda, L. Terradot, E. Marza, R. Fronzes, Filamentation of the bacterial bi-functional alcohol/aldehyde dehydrogenase AdhE is essential for substrate channeling and enzymatic regulation. *Nat Commun* **11**, 1426 (2020).
